# Demographic performance of European tree species at their hot and cold climatic edges

**DOI:** 10.1101/801084

**Authors:** Georges Kunstler, Arnaud Guyennon, Sophia Ratcliffe, Nadja Rüger, Paloma Ruiz-Benito, Dylan Z. Childs, Jonas Dahlgren, Aleksi Lehtonen, Wilfried Thuiller, Christian Wirth, Miguel A. Zavala, Roberto Salguero-Gomez

**Affiliations:** Univ. Grenoble Alpes, INRAE, LESSEM, 2 rue de la Papeterie-BP 76, F-38402 St-Martin-d’Hères, France; Department of Systematic Botany and Functional Biodiversity, University of Leipzig, Johannisallee 21-23, 04103 Leipzig, Germany; NBN Trust, 14-18 St. Mary’s Gate, Lace Market, Nottingham NG1 1PF, UK; German Centre for Integrative Biodiversity Research (iDiv) Halle-Jena-Leipzig, Deutscher Platz 5e, 04103 Leipzig, Germany; Smithsonian Tropical Research Institute, Apartado 0843-03092, Balboa, Ancón, Panama; Environmental Remote Sensing Research Group, Department of Geology, Geography and the Environment, Universidad de Alcalá, Spain; Forest Ecology and Restoration Group, Departamento de Ciencias de la Vida, Universidad de Alcala, Spain.Grupo de Ecologia y Restauracion Forestal, Departamento de Ciencias de la Vida, Universidad de Alcala, Edificio de Ciencias, Campus Universitario, 28805 Alcala de Henares, Madrid, Spain; Department of Animal & Plant Sciences, The University of Sheffield, Sheffield, UK; Swedish University of Agricultural Sciences, Umea, 90183 Sweden; Natural Resources Institute Finland (Luke), Latokartanonkaari 9 FI-00790 Helsinki Finland; Univ. Grenoble Alpes, CNRS, Univ. Savoie Mont Blanc, CNRS, LECA, Laboratoire d’Ecologie Alpine, F-38000 Grenoble, France; Max-Planck-Institute for Biogeochemistry, Hans-Knöllstr. 10, 07745 Jena, Germany; Department of Zoology, University of Oxford, 11a Mansfield Rd OX1 3SZ, Oxford, UK

**Keywords:** demography, IPM, passage time, vitale rate, climatic range edge

## Abstract

1. Species range limits are thought to result from a decline in demographic performance at range edges. However, recent studies reporting contradictory patterns in species demographic performance at their edges cast doubt on our ability to predict climate change demographic impacts. To understand these inconsistent demographic responses at the edges, we need to shift the focus from geographic to climatic edges and analyse how species responses vary with climatic constraints at the edge and species’ ecological strategy.
2. Here we parameterised integral projection models with climate and competition effects for 27 tree species using forest inventory data from over 90,000 plots across Europe. Our models estimate size-dependent climatic responses and evaluate their effects on two life trajectory metrics: lifespan and passage time – the time to grow to a large size. Then we predicted growth, survival, lifespan, and passage time at the hot and dry or cold and wet edges and compared them to their values at the species climatic centre to derive indices of demographic response at the edge. Using these indices, we investigated whether differences in species demographic response between hot and cold edges could be explained by their position along the climate gradient and functional traits related to their climate stress tolerance.
3. We found that at cold and wet edges of European tree species, growth and passage time were constrained, whereas at their hot and dry edges, survival and lifespan were constrained. Demographic constraints at the edge were stronger for species occurring in extreme conditions, i.e. in hot edges of hot-distributed species and cold edges of cold-distributed species. Species leaf nitrogen content was strongly linked to their demographic responses at the edge. In contrast, we found only weak links with wood density, leaf size, and xylem vulnerability to embolism.
4. Synthesis. Our study presents a more complicated picture than previously thought with demographic responses that differ between hot and cold edges. Predictions of climate change impacts should be refined to include edge and species characteristics.

## Introduction

In the face of climate change there are increasing concerns about the future redistribution of plant species ranges (Zimmermann et al., 2013). Range shifts are thought to be directly related to changes in population dynamics. The classical view of the link between population dynamics and species ranges comes from a long-standing hypothesis in biogeography known as the ‘abundant-centre hypothesis’ (hereafter ACH, Brown, 1984; Pironon et al., 2017), which proposes that demographic performance decline at the range edge results in a decrease in abundance, occupancy and genetic diversity (Pironon et al., 2017). This is directly related to the hypothesis that at equilibrium, a species’ range edge should occur where the mean population growth rate (*λ*) drops below one (*λ <* 1) due to changes in one or more vital rates (*i.e.* survival, growth, or reproduction) (Case, Holt, McPeek, & Keitt, 2005; Holt & Keitt, 2005).

Understanding the demographic pathways of population response at range edges is crucial for forecasting climate change impacts. However, existing studies comparing population growth rates or vital rates in the periphery *vs.* the centre of species geographic range provide weak support for the ACH (Pironon et al., 2017). Transplant experiments have shown a decline in population growth rate or vital rates beyond the geographic edge but not necessarily right at the edge (Hargreaves, Samis, & Eckert, 2014; Lee-Yaw et al., 2016). Similarly, model-based analyses of natural population monitoring data have found no clear evidence of a decrease in demographic performance at the geographic edge (Csergo et al., 2017; Purves, 2009).

Recent reviews have highlighted the difficulties of synthesising existing results because most studies explored performance of geographically peripheral populations without a clear understanding of the local climatic or environmental constrains (Pironon et al., 2017; Vilà-Cabrera, Premoli, & Jump, 2019). Changes in demographic performance are, however, likely to vary depending on the type of biophysical constraints at the edge (Gaston, 2009) and therefore, demographic performance at the edge should be analysed in relation to the local main climatic contraints (the “central-marginal” hypothesis in Pironon et al., 2017). Firstly, demographic constraints could differ between drought- and cold-limited edges because tolerance to different abiotic stresses requires different adaptative strategies (Niinemets & Valladares, 2006) resulting in different vital rates being constraint at these edges (Gaston, 2009; Hargreaves et al., 2014). Secondly, it has been proposed that biotic interactions (*e.g.* competition) could be key constraints of demographic performance at the edge and that this effect would be stronger for edges in productive environments than in unproductive environments. However, support for this hypothesis is limited (see Hargreaves et al., 2014; Cahill et al., 2014; Louthan, Doak, & Angert, 2015). Thirdly, constraints on the demographic performance at a climatic edge are likely to vary with species’ physiological strategy (Anderegg, Anderegg, Kerr, & Trugman, 2019). These physiological differences can be captured by species’ climatic optimum and by functional traits related to species physiological climate response, such as wood (Chave et al., 2009) or leaf characteristics (Wright et al., 2017). Finally, an additional difficulty arises in long-lived organisms such as trees because the response of their vital rates to climatic constraints at the edge might vary depending on the size of the individual (Tredennick, Teller, Adler, Hooker, & Ellner, 2018). This size-dependent response to climate can be crucial for size-structured populations (De Roos, Persson, & McCauley, 2003; Tredennick et al., 2018) and can affect the population performance at the edge. We thus need to analyse the performance at the edge with size-structured models translating size-dependent climatic responses and the demographic compensation effect they may occur between size or vital rates into life trajectory metrics.

Here, we explored these questions in European forests, which play a crucial role for multiple ecosystem services such as sheltering a significant proportion of biodiversity and carbon stocks and contributing to the livelihoods of local populations (van der Plas et al., 2018). We used size-structured models fitted to forest inventory data documenting survival and growth of more than one million adult trees across the continent covering Mediterranean, temperate and boreal biomes. Firstly, we fitted survival and growth models for 27 species to capture size-dependent climate and competition responses of these vital rates. Secondly, we built size-structured population models using integral projection models (IPM) (Ellner, Childs, & Rees, 2016) to evaluate how size-dependent responses to climatic constraints at the edge translate into two life trajectory metrics - mean lifespan and passage time (time to grow from small to large size). We then used these models to compare species’ predicted demographic performance at the hot and dry or cold and wet climatic edges with their performance at the climatic centre. Using these metrics we tested the following hypotheses: (1) vital rates and IPM-derived performance metrics are reduced at the climatic edge compared to the climatic centre but the demographic metrics affected differ between cold and hot edges; (2) the decline in demographic performance at the climatic edges is stronger in the presence of local competition than without; and (3) demographic performance at the climatic edge depends on species’ position along the climate gradient and functional traits related to species’ climate stresses tolerance (testing the effect of wood density, leaf economic spectrum traits, leaf size, and xylem vulnerability to embolism).

## Materials and Methods

In this section we present: (1) the development of climate-dependent IPMs based on growth and survival models and the data used to fit them; (2) the development of species distribution models used to select climatic edges corresponding to a species distribution limits; (3) the derivation of metrics of demographic performance at the climatic edge *vs.* the climatic center of the species distribution from the IPMs; and (4) the methodology to test our three hypotheses.

### Forest inventory

We used the European forest inventory (NFI) data compiled in the FunDivEUROPE project (Baeten et al., 2013; Ratcliffe et al., 2015). The data covers 91,528 plots and more than one million trees in Spain, France, Germany, Sweden and Finland. NFIs record information on individual trees in each plot, including species identity, diameter at breast height (dbh), and status (alive, dead, harvested or ingrowth). Plot design varies between countries but generally plots are circular with variable radii depending on tree size (see Supporting Information). The minimum dbh of trees included in the dataset was 10 cm and plots were remeasured over time allowing estimations of individual growth and survival. The time between two survey varied from 4 to 16 years. Only the French NFI is based on a single measurement but provides a measurement of radial growth from cores (over 5 years) and an estimation of time since death. We selected species with > 2,000 individuals and > 500 plots, to ensure a good coverage of their range, growth, and survival. We excluded exotic species for which the distribution is mainly controlled by plantation operations. For the demographic analyses, we also excluded all plots with records of harvesting operations or disturbances between the two surveys, which would otherwise influence our estimation of local competition.

### Climate variables

We used two bioclimatic variables known to control tree demography (Kunstler et al., 2011): (1) the sum of degree days above 5.5 °C (*sgdd*), and (2) the water availability index (*wai*). *sgdd* is the cumulative day-by-day sum of the number of degrees > 5.5 °C and is related to the mean annual temperature and the length of the growing season. It was extracted from E-OBS, a high resolution (1 km^2^) downscaled climate data-set (Moreno & Hasenauer, 2016) for the years between the two surveys plus two years before the first survey. In preliminary analyses we also explored the number of frost days but it was too correlated with *sgdd* to be included in the models. *wai* was computed using precipitation (*P*, extracted from E-OBS) and potential evapotranspiration (*PET*) from the Climatic Research Unit (Harris, Jones, Osborn, & Lister, 2014) data-set, as (*P − PET*)/*PET* (see Ratcliffe et al., 2017) and is related to the water availability. We also explored other water stress indices but they did not improve the demographic models so we decided to use *wai*.

### Integral projection models

An IPM predicts the size distribution, *n*(*z^t^*, *t* + 1), of a population at time *t* + 1 from its size distribution at *t*, *n*(*z*, *t*), where *z* the size at *t* and *z^t^* the size at *t* + 1, based on the following equation (Easterling, Ellner, & Dixon, 2000; Ellner et al., 2016):

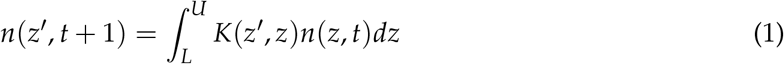

The kernel *K*(*z^t^*, *z*) can be split into the survival and growth kernel (*P*(*z^t^*, *z*)) and the fecundity kernel (*F*(*z^t^*, *z*)), as follow *K*(*z^t^*, *z*) = *P*(*z^t^*, *z*) + *F*(*z^t^*, *z*). *P*(*z^t^*, *z*) is defined as *P*(*z^t^*, *z*) = *s*(*z*)*G*(*z^t^*, *z*) and represents the probability that an individual of size *z* survives between *t* and *t* + 1 and reaches the size *z^t^*. The size of the individuals *z* can range between *L* and *U*. NFI data do not provide direct information on tree fecundity, thus our models describe the fate of a cohort (a cohort IPM for individuals with dbh *>*= 10 cm) by focusing only on *P*(*z^t^*, *z*). Even without covering the full life cycle, cohort IPMs are useful to estimate demographic performance because they allow to predict life trajectory metrics accounting for size-dependent climate responses and compensatory effect between vital rates.

For each of the 27 species, we fitted growth and survival functions depending on tree size, the two climatic variables (*sggd* and *wai*) and local competition estimated as the sum of basal area of competitors (following Kunstler et al., 2011). The shape of the climatic response curves and the type of interaction between climate and tree size and climate and competition (which represents a size-dependent response) can have a large impact on vital rates predictions and IPM derived life trajectory metrics. To account for such uncertainties, we re-sampled 100 times 70% of the data to fit the growth and survival models and select the best type of climatic response curves and interactions based on the Akaike information criteria (*i.e.*, lowest AIC) (Burnham & Anderson, 2002). Because there were fewer plots in extreme climatic conditions, we re-sampled the data with a higher probability of sampling plots in extreme climatic conditions for the given species (see details in Supporting Information). Then we used the remaining 30% of the data to evaluate the goodness of fit of the growth and survival models. Goodness of fit and response curves of growth and survival models are presented in the Supporting Information (Figs 4 to 13).

#### Growth model

After preliminary exploration, we selected two alternative shapes of the climatic response curves: asymptotic or quadratic polynomial corresponding respectively to the equation 2 and the equation 3. These equations are flexible and allow for increasing, decreasing, bell or U-shape responses. These two equations also allowed to represent two alternative biological models: (i) either all species have their optimum at high water availability and sum of degree days; or (ii) species have bell-shaped climate response curves with different optima along the climatic variables.

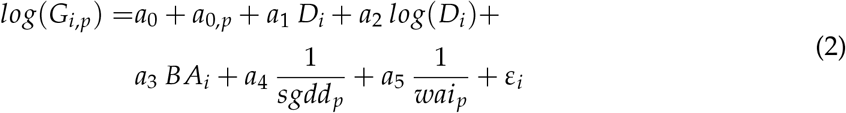

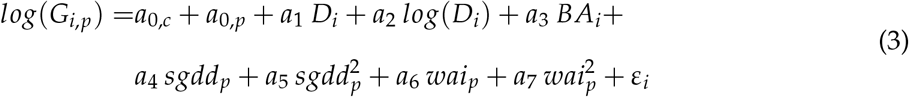

Where *G_i_*_,*p*_ is the annual diameter growth of tree *i* in plot *p*, *D_i_* is the dbh of tree *i*, *BA_i_* is the sum of basal area of local competitors of tree *i* per ha (sum basal area of both conspecific and heterospecific trees in the plot in a single local competition index), *sgdd_p_* is the sum of growing degree days, *wai_p_* is the water aridity index, *a*_0_ to *a*_7_ are estimated parameters, and *a*_0,*p*_ is a normal random plot effect accounting for unexplained variation at the plot level. The intercept *a*_0,*c*_ is country specific to account for differences in sampling protocol between the NFIs and *ε_i_* is the unexplained tree level variability following a normal distribution. We also tested models with interactions between the climatic variables - 1/*sgdd_p_* and 1/*wai_p_* for model (2) and *sgdd_p_* and *wai_p_* for model (3)) - and size (*D_i_* and *log*(*D_i_*)) and the climatic variables and competition. We fitted the models in R-cran separately for each species (R Core Team, 2019) using the ‘lmer’ function (“lme4” package, Bates, Mächler, Bolker, & Walker, 2015).

#### Survival model

Survival models were fitted with a generalised linear model with a binomial error. The predictors and interactions explored were the same as in the growth model. To account for variable survey times between plots we used the complementary log-log link with an offset representing the number of years between the two surveys (*y_p_*) (Morris, Vesk, & McCarthy, 2013). We fitted the model in R-cran using the ‘glm’ function. We did not include a random plot intercept because in most plots no individuals died between the surveys, making the estimation of the random plot effect challenging.

#### Tree harvesting

Although we excluded plots with evidence of harvesting between the two surveys to fit the survival functions, most European forests are subject to management, which has a strong impact on population dynamics (Schelhaas et al., 2018). Preferential harvesting of dying or damaged trees before their death probably results in an underestimation of the natural mortality rate. To make sensible predictions with our IPMs it was necessary to incorporate a harvesting rate to prevent an overestimation of tree lifespan. We set the individual tree harvesting rate, as the mean harvesting annual probability across all species and countries. The estimate was 0.5% per year. We did not model size and climate dependence of the harvesting rate, as we focused on climatic and not anthropogenic constraints on tree demography.

### Prediction of demographic metrics at the climatic edges and centre of species range

#### Species distribution

To identify the climatic edge of a species range, a simple representation of its distribution in climate space is necessary. Across Europe, there is a strong correlation between *sgdd* and *wai*, and so we described species ranges along a single climatic axis corresponding to the first axis (PC1) of the PCA of *sgdd* and *wai* (Supporting Information, Fig. 3). Species showed a clear segregation along this climatic axis in Europe (Fig. 1). Based on the coordinates on PC1 of the plots where the species was present, we identified the median climate as their median value of PC1 (which we used as an index of species position along the climate gradient), the hot and dry edge (hereafter hot edge) and the cold and wet edge (hereafter cold edge), respectively, as their 5% and 95% quantiles. These quantiles represent two extreme climatic conditions experienced by the species. By focusing on climatically marginal populations, our approach differs from most tests of the ACH reviewed in Pironon et al. (2017) that studied populations at the periphery of the species geographic range.

**Figure 1:**
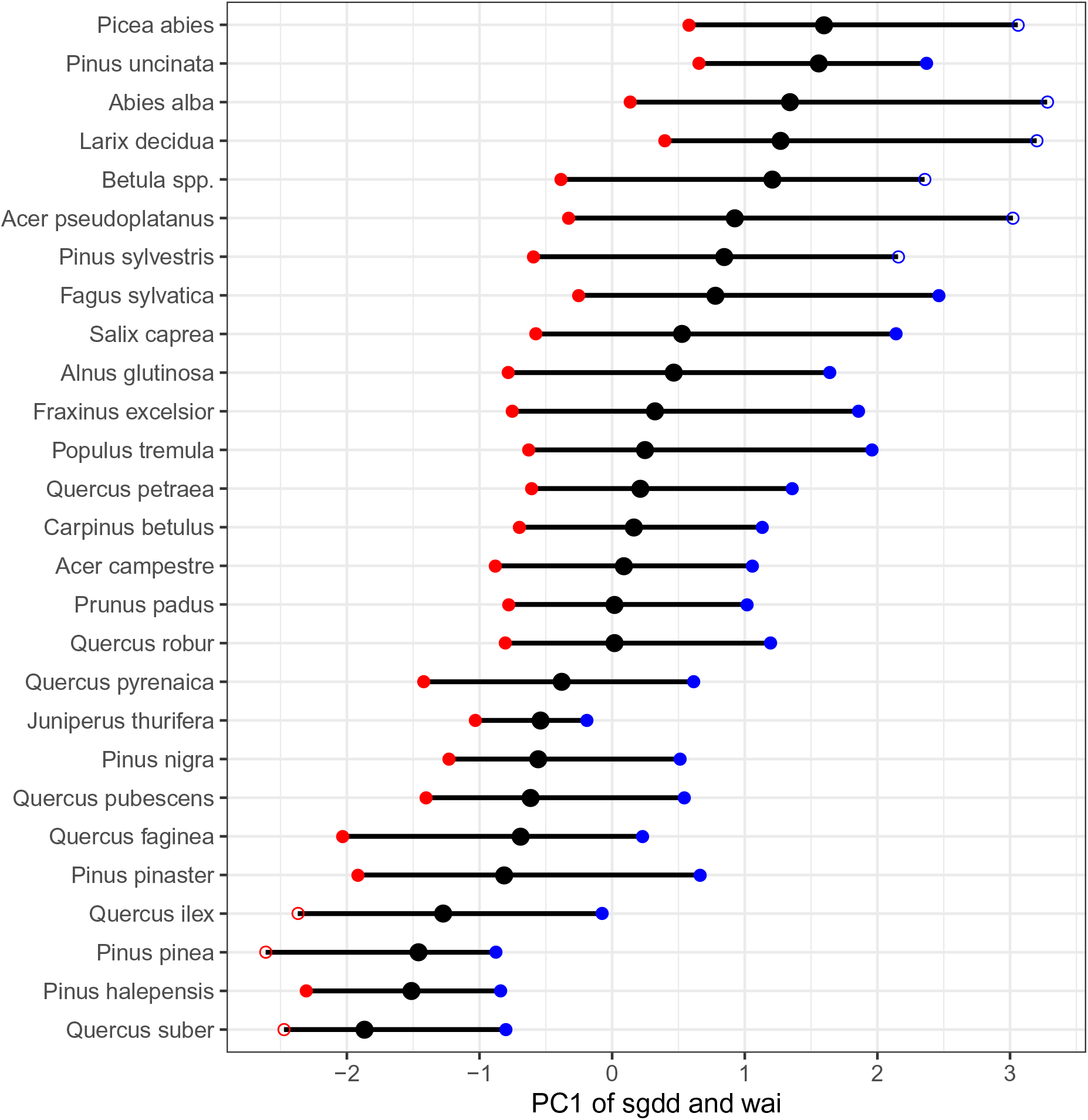
Species distribution along the first axis of the PCA of the two climatic variables *sgdd* and *wai*. The median of the species distribution along this axis is represented by a black circle and the hot and dry edge and the cold and wet edge by red and blue circle respectively. Filled circles represent edges with a clear drop of the probability of presence as predicted by species distribution models that were selected for the analysis.

To evaluate which species’ edges corresponded to an actual limit in the species distribution and not just to limits in data coverage, we fitted species distribution models with BIOMOD2 (Thuiller, Lafourcade, Engler, & Araújo, 2009) using presence/absence data covering all Europe (Mauri, Strona, & San-Miguel-Ayanz, 2017) (see Supporting Information). For comparison of the demographic performance at the edge *vs.* the centre of the distribution, we retained only the edges where the SDM predicted at least a 10% drop in the probability of presence of the species (Fig. 1).

#### Demographic metrics

To evaluate how individual tree performance varied between the species’ median climate and the climatic edges, we derived four metrics representing key dimensions of population performance. The first two metrics were related to individual vital rates, and were defined by the growth and survival of 15 cm dbh individuals. We focused on small individuals because of their large effect on population dynamics (Grubb, 1977). The last two metrics were life trajectory metrics integrating the vital rates and size-dependent responses to climate in the IPM, and were defined by the mean lifespan of a 10 cm dbh individual and the passage time of a 10 cm dbh individual to 60 cm. The details of the numerical methods used to compute lifespan and passage time from the IPM are provided in the Supporting Information. Model diagnostics showed that our numerical approach was not sensitive to the number of size bins retained for the IPM (*i.e.* # bins > 800, see Fig. 14 in Supporting Information).

We predicted the four demographic metrics at the centre and the hot and cold climatic edges of the species using their positions on the climatic axis. The median, and 5% and 95% quantiles on the PC1 correspond to the projection of a unique combination of *sggd* and *wai* for which we predicted the metrics. We integrated uncertainty into our estimates by deriving each demographic metric for all 100 re-sampled growth and survival models (see above). Because competitive interactions may also be important in controlling species demography at the edge of the range (Louthan et al., 2015), we made these predictions either without local competition (by setting *BA* to 0) or with a high level of local competition (by setting *BA* to 30*m*^2^*ha^−^*^1^, corresponding to a closed forest).

#### Analysis of the relative demographic performance at the climatic edges

For each demographic metric (*m*) we computed the relative difference in the metric at the edge (hot or cold) *vs*. the centre as: 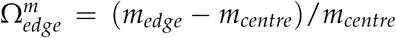. We integrated uncertainty by deriving estimates of 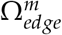 for each of the 100 re-sampled growth and survival models. Then we used 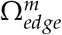 to evaluate our three hypotheses.

Firstly, for each metric, we tested whether species demographic performance declined at the climatic edge compared to the climate centre (hypothesis 1) by fitting a mixed model to test whether *m* was function of the range position type (edge *vs.* centre) using the function *lmer* in *lme4*. We included a random species effect to account for the non-independence of the 100 re-sampled estimates per species. We ran this analysis separately for hot and cold edges to see how demographic responses differed between them. Secondly, we tested whether the effects were different without or with competition (hypothesis 2).

Thirdly, we explored whether 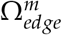 was dependent on species median climate and functional traits related to species’ climatic response (hypothesis 3). We used Phylogenetic generalised least squares (PGLS) regression using a phylogeny extracted from Zanne et al. (2014) to account for phylogenetic dependence between species. We accounted for the uncertainty in the demographic response by including a weight proportional to the inverse of the variance of 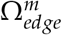 (estimated over the 100 re-sampled growth and survival models). The PGLS regression with maximum likelihood estimation of Pagel’s lambda (a measure of the phylogenetic signal ranging between 0 and 1) did not always converged. In those cases we fitted a PGLS model with a Brownian model (Pagel’s lambda set at 1). We retained only the regressions that were both significant (after a Bonferroni correction to account for multiple comparisons) and had a non-negligible magnitude of the effect (Camp, Seavy, Gorresen, & Reynolds, 2008). The magnitude of the effect was considered negligible when the confidence interval of the effect size intercepted the interval -0.10 and 0.10 (Camp et al., 2008). Effect sizes were computed as the standardised slope (Schielzeth, 2010).

To test the link between 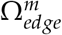 and species median climate, we ran the PGLS regression between 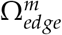 and the species median position on PC1. To test the links between 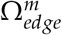 and functional traits, we ran the same type of PGLS regressions with four functional traits that are known to influence tree response to climate. We selected the following traits: (i) wood density, because of its links with drought and temperature response (Chave et al., 2009; Stahl, Reu, & Wirth, 2014); (ii) the leaf economic spectrum (LES) because species at the conservative end of the spectrum are thought to be more tolerant to extreme climate (Reich, 2014); (iii) leaf size, because of its links with water stress and frost response (Wright et al., 2017); and (iv) xylem vulnerability to embolism measured by the water potential leading to 50% loss of xylem conductivity, Ψ_50_, because of its link with drought-induced mortality (Anderegg et al., 2016). LES is based on the covariance of specific leaf area, leaf lifespan, and leaf nitrogen per mass (Wright et al., 2004). We used leaf nitrogen per mass (*N_mass_*), as it was the LES trait with the best coverage across our species. Trait data were sourced from open databases (Chave et al., 2009; Choat et al., 2012; Maire et al., 2015; Wright et al., 2017, 2004).

## Results

### Growth and survival size-dependent responses to climate

For most species the growth and survival models showed evidence of interactions between climate and tree size and for a smaller subset of species also between climate and competition (see Tables 2 and 3 Supporting Information). This indicates that size-dependent climatic responses were common. Model selection over the 100 re-sampled data showed that for 23 species out of 27 the most frequently selected growth model included interactions between climate variables and tree size (see Table 2 in Supporting Information). Selection of the best survival model was more variable between the 100 data re-sampling than for the growth models. For 17 species out of 27 the most frequently selected survival models included interactions between climatic variables and tree size (see Table 3 Supporting Information). For both growth and survival several species also showed evidence of interactions between climate variables and competition (respectively 12 and 11 species out of 27, see Tables 2 and 3 Supporting Information).

### Demographic responses differ between edge types and metrics

Across the 27 species, we found evidence of a significant decrease in growth and increase in passage time (longer time needed to grow from 10 to 60 cm) at the cold edge in comparison with the median climate but no effect at the hot edge (Fig. 2). In contrast, at the hot edge, we found evidence of a significant decrease in both tree survival and lifespan (Fig. 2). This is consistent with the hypothesis that at least one metric will decline in performance at the edge, and that different metrics are affected depending on the edge type. In contrast, we found that lifespan was significantly longer at the cold edge than at the median climate (Fig. 2). Generally, these patterns were unaffected by local competition (Fig. 3). It is, however, important to note that the relative decrease in survival at the hot edge and the increase of passage time at the cold edge became non-significant at high levels of competition (Fig. 3).

**Figure 2:**
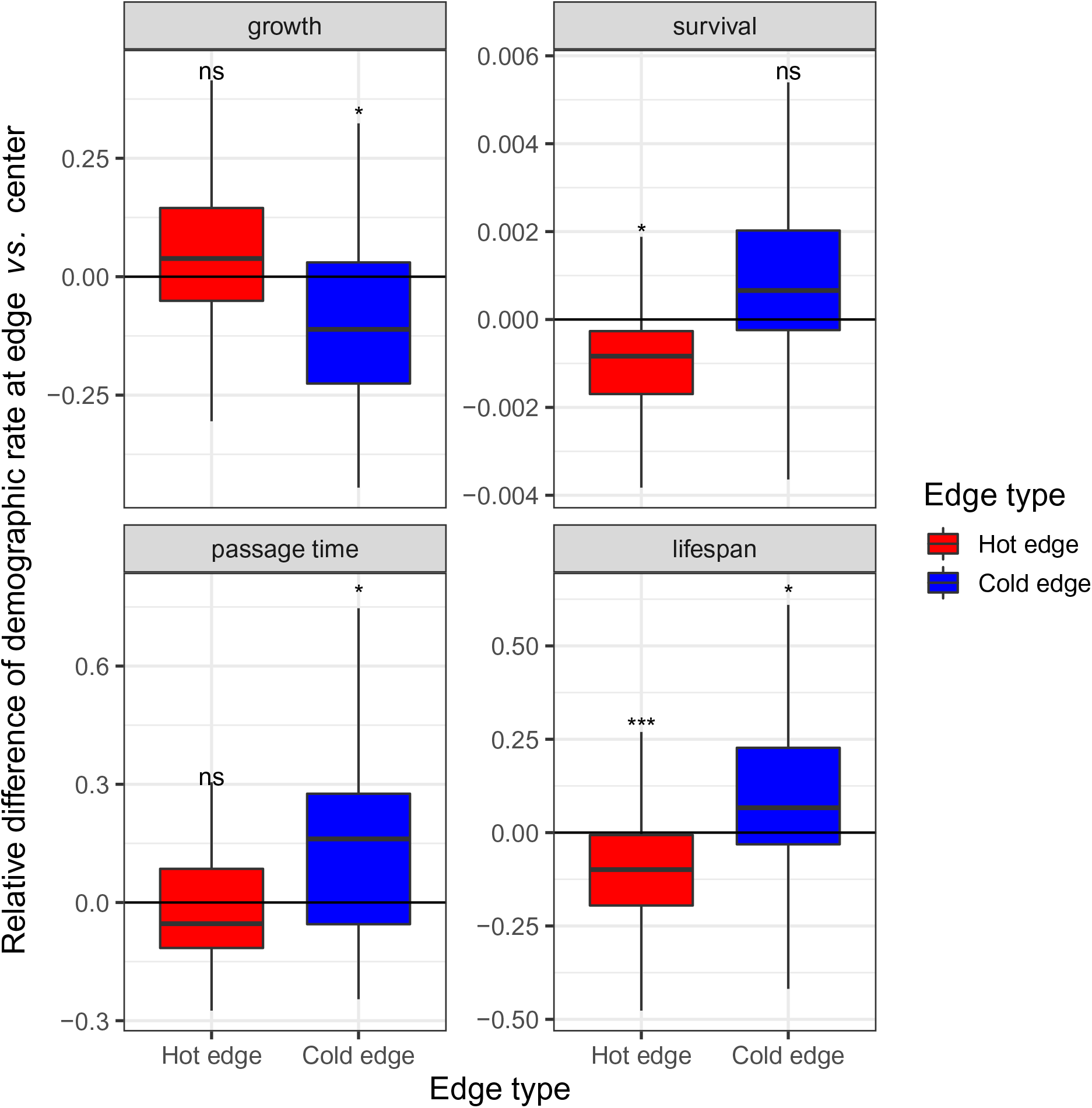
Differences in the demographic metrics at climatic edge *vs.* the median climate of the species distribution. The box-plots represent the relative difference of the demographic metrics between the climatic edge and the median climate computed over the 100 data resampling and the 27 species for the four demographic metrics (annual diameter growth and survival for an individual 15cm in diameter, passage time from 10cm in diameter to 60cm in diameter and lifespan of tree 15cm in diameter) and the two edge types (hot in red, cold in blue). The p value of the test for the difference in each demographic metric between the edge and the median climate is presented at the top of the box-plot (ns : non significant, * : p value < 0.05, ** : p value < 0.01, *** : p value < 0.001). The p value was computed with a mixed model with species as a random effect (see Methods for details).

**Figure 3:**
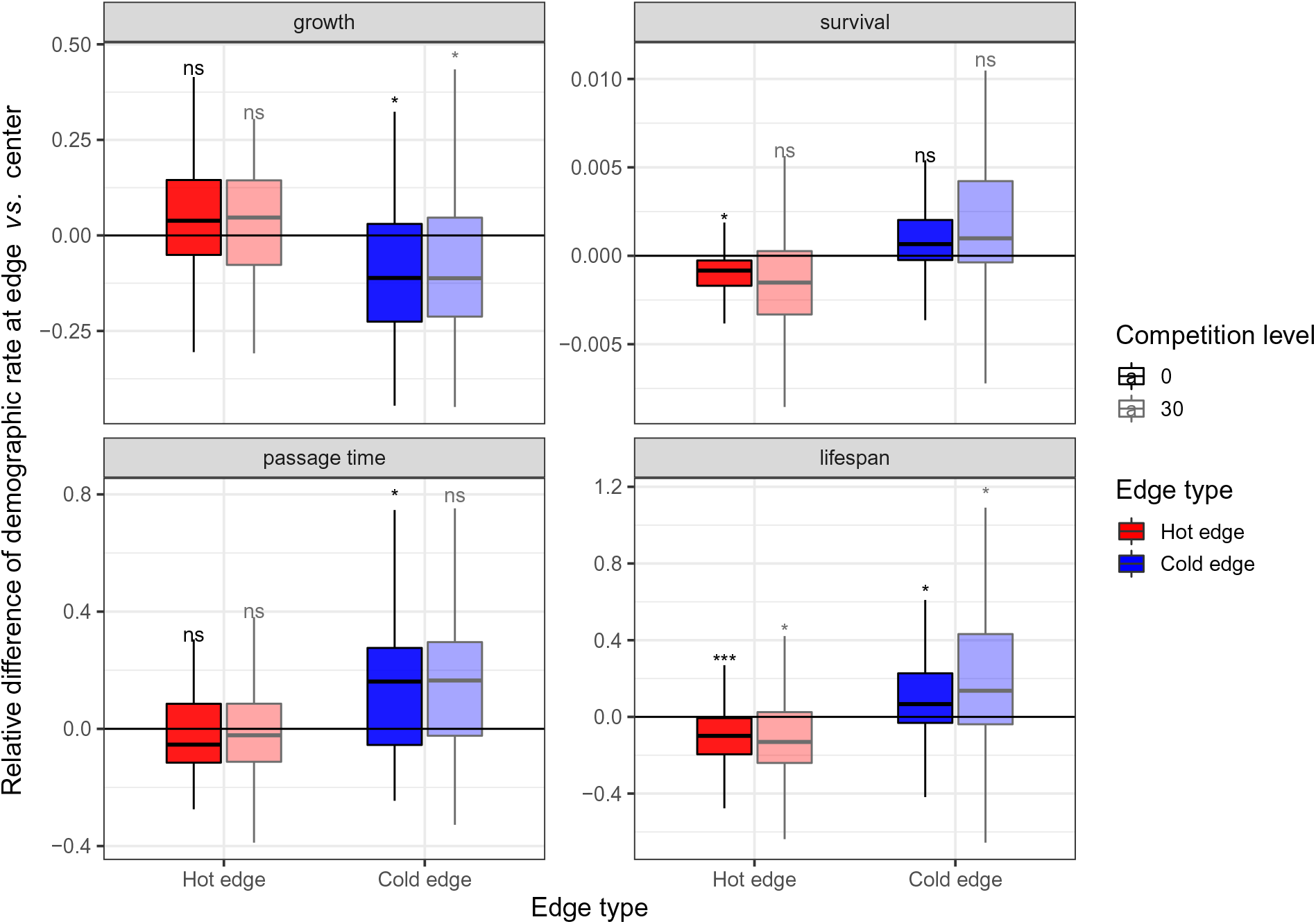
Differences in the demographic metrics at climatic edge *vs.* the median climate of the species distribution without and with a high level of competition. The box-plots represent the relative difference the demographic metrics between the climatic edge and the median climate over the 100 data resampling and the 27 species for the four demographic metrics (annual diameter growth and survival for an individual 15cm in diameter, passage time from 10cm in diameter to 60cm in diameter and lifespan of tree 15cm in diameter), the two edge types (hot in red, cold in blue), and the two levels of competition (without competition: basal area of competitors, *BA* = 0, no transparency, with a high level of competition: basal area of competitors, *BA* = 30*m*^2^ *ha^−^*^1^ color transparency). The p value of the test for the difference in each demographic metric between the edge and the median climate is presented at the top of the box-plot (ns : non significant, * : *p*-value < 0.05, ** : *p*-value < 0.01, *** : *p*-value < 0.001). The *p*-value was computed with a mixed model with species as a random effect (see Methods for details)

Despite the overall demographic response at the edge, there were large variations between species. For each metric and edge type we found species showing a decrease and species showing an increase in performance (Supporting Information; Figs 16 to 19).

### Demographic responses vary with species median climate

Growth response at the hot and cold edges was related to the median climate of the species; species associated with hot climates were more constrained at their hot edge while species associated with cold climates were more constrained at their cold edge. This result is depicted in Fig. 4 by a positive relationship between the median climate of the species and 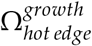 and a negative relationship with 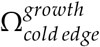. The same pattern is visible for passage time, but in the opposite direction, because passage time is longer when growth is slower (Fig. 4). The responses of 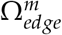 for survival and lifespan were much weaker or null. We found a negative relationship for 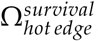, which was largely related to a few extreme species, and no effect for lifespan (Fig. 4).

**Figure 4:**
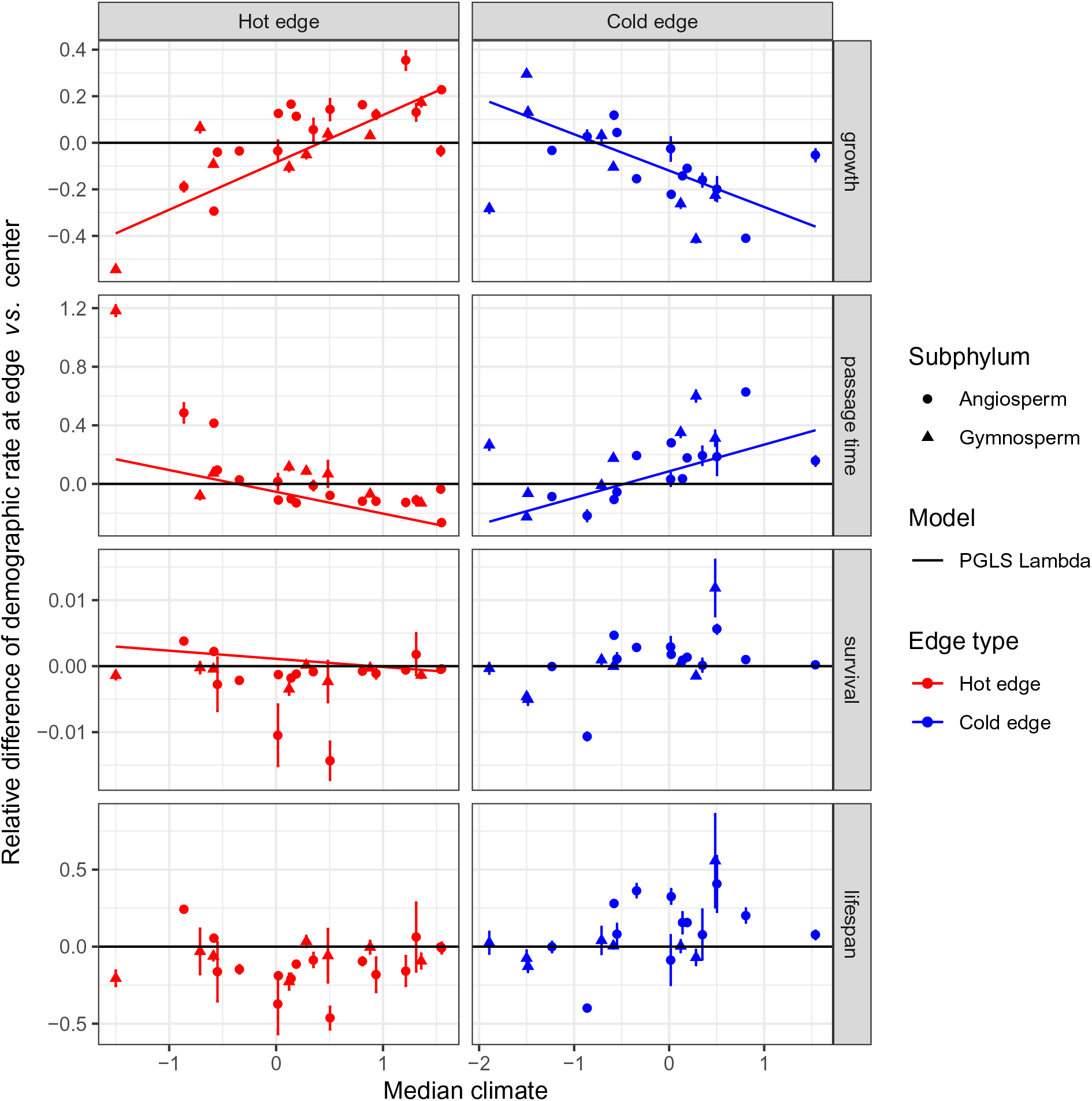
Changes in demographic responses at the edge in function of species median climate. Species demographic response at the edge - measured as the relative differences of the demographic metrics at climatic edge *vs.* the median climate of the species distribution - in function of the median position of the species on the first axis of the climate PCA. For each species the mean (point) and the 95% quantiles (error bar) of the demographic response over the 100 data resampling is represented for both the hot (red) and the cold (blue) edges. Phylogenetic generalised least squares (PGLS Lambda) regressions are represented only for significant relationship with a non negligible magnitude of the effect. Gymnosperm and angiosperm species are represented with different symbols.

### Weak links between demographic response and species traits

*N_mass_* had the strongest relationship with 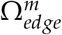 of all the traits we tested. At the hot edge, species with high *N_mass_* experienced a stronger decrease in their survival and lifespan than species with a low *N_mass_* (Fig. 5). In contrast, at the cold edge, species with low *N_mass_* experienced a stronger decrease in their survival and lifespan than species with high *N_mass_* (Fig. 5). In addition, species with high *N_mass_* had less limitation of their growth at the hot edge than species with low *N_mass_* (Fig. 5). In contrast, species with high *N_mass_* had stronger limitation of their growth at the cold edge (Fig. 5).

**Figure 5:**
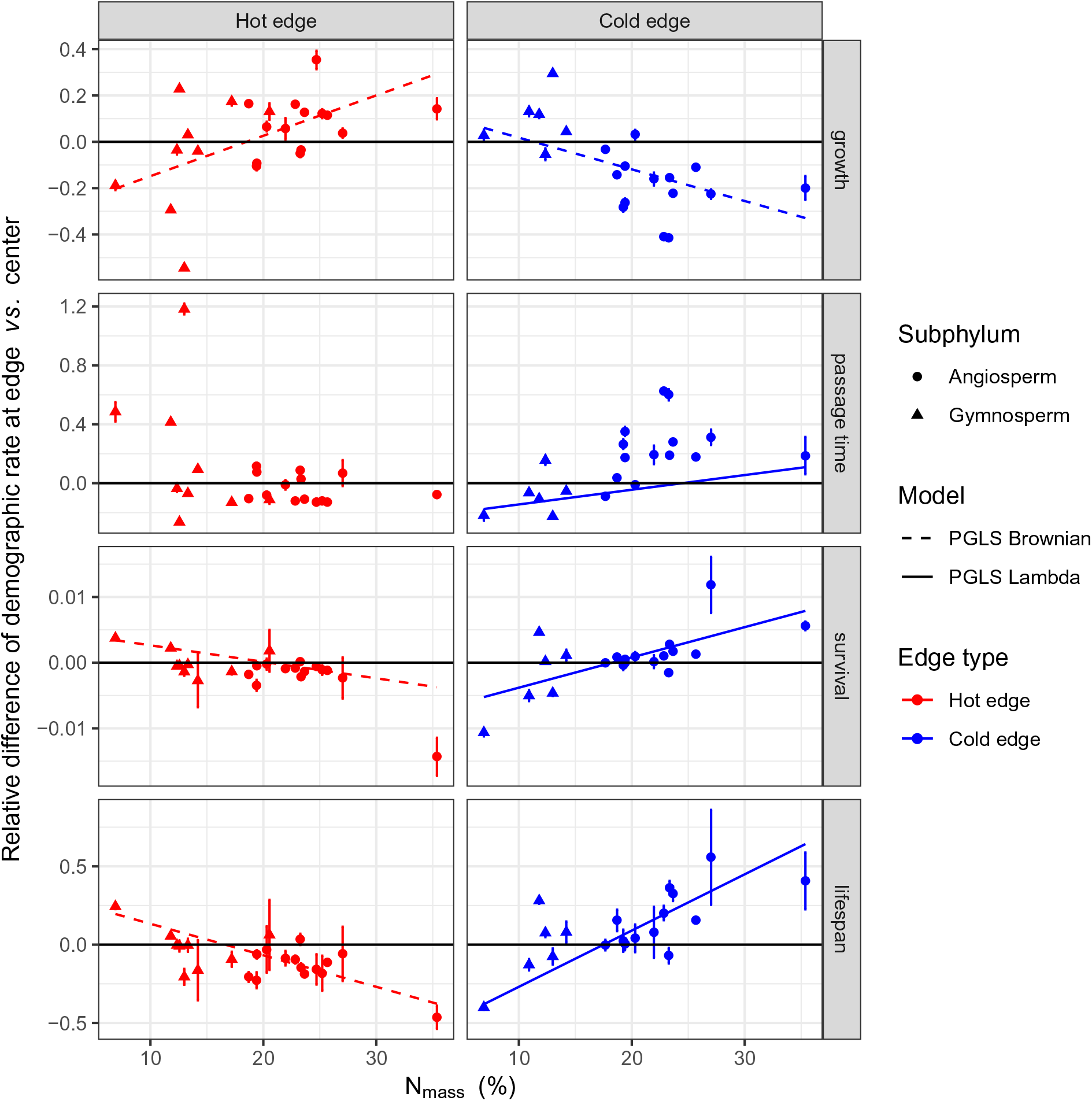
Changes in demographic responses at the edge in function of species leaf N per mass. Species demographic response at the edge - measured as the relative differences in the demographic metric at climatic edge *vs.* the median climate of the species distribution - as a function of species leaf nitrogen per mass. For each species the mean (point) and 95% quantiles (error bar) of the demographic response over the 100 data resampling is represented for both the hot (red) and the cold (blue) edges. Phylogenetic generalised least squares (PGLS) regressions are represented only for significant relationship with a non negligible magnitude of the effect (see details in caption of Fig. 4).

Relationships between 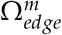 and wood density, leaf size and xylem vulnerability to embolism (Ψ_50_) were generally weak (Supporting Information, Figs 21 to 23). Most of these relationships were driven by only a few species (Supplementary Information, Figs 21 22 23). Species with small leaf area had better survival and lifespan at the hot edge and better passage time at the cold edge than large leafed species (Supplementary Information, Fig. 23). Species with high Ψ_50_ experienced a stronger decrease in their growth at the hot edge than species with low Ψ_50_ (Supplementary Information, Fig. 22).

## Discussion

Our analysis based on pan-European forest inventory data and integral projection models of 27 tree species, found weak support for the ACH prediction that demographic performance is lower at the climatic edge than at the centre of the species range. Instead, decline in demographic performance was strikingly different between the cold and the hot edges. At cold and wet edges, growth and passage time were constrained, whereas at hot and dry edges, survival and lifespan were constrained. Beyond these general patterns, we found important variability between species in their demographic performance at the edge, which was partially explained by species’ median climate and traits.

### Different demographic responses at the hot and the cold edge

We found mixed support for the ACH; not all the demographic metrics were limited at the two edges and patterns were variable between species. This is consistent with observational studies that found limited evidence of a relationship between species demography and their distribution. For instance, both Thuiller et al. (2014) and Csergo et al. (2017) found limited correlation between plants demographic performance and probability of presence. In addition, Purves (2009) reported mixed evidence of a decrease in demographic performance at the south and north edges of North American tree species.

Growth and passage time were constrained at the cold edge in comparison with the centre of the species climatic range. This is consistent with studies on North American tree species, that found a decrease in growth at the cold edge in adult trees (Purves, 2009) and juveniles (Ettinger & HilleRisLambers, 2017; Putnam & Reich, 2017). In contrast with the ACH, we found a tendency for a slightly faster growth at the hot edge than at the centre, which has also been reported in North American trees (Ettinger & HilleRisLambers, 2017; Purves, 2009; Putnam & Reich, 2017). Interestingly, studies on *Fagus sylvatica* radial growth in Europe found a higher drought resistance at the hot edge than at the core of the range (Cavin & Jump, 2017).

At the hot and dry edge, tree survival and lifespan were lower than at the centre of the climatic range. The same decrease of survival at the hot edge was also found by Archambeau et al. (2020) for *Fagus sylvatica* and *Pinus sylvestris* in Europe. In contrast, Purves (2009) found no such decrease in survival at the hot edge of eastern North American species. This difference could be explained by the fact that the hot edge of most European species corresponds to both a hot and a dry climate, whereas in eastern America the hot edge is less constrained by drought (Zhu, Woodall, Ghosh, Gelfand, & Clark, 2014). We found that lifespan was longer at the cold edge than at the centre of the distribution, which contradicts the classical view that survival is constrained in cold climates and the results of Purves (2009). Given that tree diameter growth is constrained at the cold edge, this longer lifespan could be explained by a tradeoff between tree growth rate and tree longevity (see Black, Colbert, & Pederson, 2008; Di Filippo et al., 2015) and the observation that survival rate correlates negatively with site productivity (Stephenson et al., 2011).

We found strong evidence of size-dependence of growth and survival responses to climatic constraints. Our results agree with previous studies which found that tree growth or survival responses to climate varied with ontogeny (Canham & Murphy, 2017; Trouillier et al., 2019). For instance, Canham & Murphy (2017) found a displacement of the climatic optimum of growth and survival between seedlings, saplings, and canopy trees. These size-dependent climatic responses, however, did not strongly influence the life trajectory metrics derived with IPMs as the response of lifespan at the edge was closely connected to the survival of a 15 cm dbh tree and the passage time was closely related to the growth of a 15 cm dbh tree. This means that these size-dependent responses were either of small magnitude or led to few compensation effects between size classes. Tredennick et al. (2018) also found that the size-dependence of vital rates responses to exogenous environmental fluctuations had limited effect on the population growth rate of perennial plant species.

### Lack of competition effect

Numerous studies have proposed that competitive interactions could be crucial in setting demographic limits, particularly when site productive is high (see Hargreaves et al., 2014; Alexander, Diez, Hart, & Levine, 2016; Ettinger & HilleRisLambers, 2017; HilleRisLambers, Harsch, Ettinger, Ford, & Theobald, 2013; Louthan et al., 2015). In our analyses, we explored the effect of competition by comparing the relative demographic performance at the edge in comparison with the centre (Ω) without local competition or with a high level of local competition. Despite the strong direct effects of competition on both growth and survival and interactions between competition and climate (see the variables importance reported in Supplementary Information, Tables 4 and 5), the relative demographic responses at the edges *vs.* the centre (measured by Ω) were not strongly influenced by the degree of local competition. Competition is thus a strong determinant of demographic rates but its effect is not stronger at the climatic edge than at the climatic centre (rejecting hypothesis 2). Rather competition blurs demographic constraints at the edge. Indeed, limitations of survival at the hot edge and passage time at the cold edge were significant without competition but not with a high level of competition.

Three main reasons could explain the lack of competition effect on the demographic response at the edges in our study. Firstly, properly estimating competition effect with observational data is notoriously difficult (Tuck, Porter, Rees, & Turnbull, 2018). Secondly, we did not differentiate between intra- and inter-specific competition, whereas inter-specific competition might have the strongest impact at the edge (Alexander et al., 2016). Thirdly, as our cohort IPMs do not cover the full life cycle it was not possible to evaluate whether competitive exclusion - the final effect of competition (Chesson, 2018) - occurs at the edge.

### Strong effect of species median climate on growth response at the edge

We found that the hotter the centre of the species range, the greater were the constraints on growth and passage time at its hot edge. The same pattern was found with the cold edge and the species median climate proximity to cold extreme. This is in agreement with the general observation that, in Europe, vegetation productivity in Europe is at its maximum in temperate climates where both drought and cold stress are limited (Jung et al., 2007).

### Weak trait effect on species demographic response at the edge

Part of the variation in the demographic response at the edge between species was related to *N_mass_*, a key dimension of the leaf economic spectrum. An important difficulty in the interpretation of these results is that our understanding of the link between leaf economic traits and climate is limited. Multiple mechanisms, some of them contradictory, have been proposed to explain the link between leaf N and climate. For instance, it is generally considered that species with low *N_mass_* have a more conservative strategy of resource use and perform better in stressful conditions than species with high *N_mass_* (Reich, 2014). In agreement with this finding, we found that species with low *N_mass_* had a better survival and lifespan at the hot edge. In contrast, high leaf N has been linked with photosynthesis tolerance to drought and low temperatures because of higher enzyme activities (Reich & Oleksyn, 2004; Wright, Reich, & Westoby, 2003). Consistent with this mechanism, we found that species with high *N_mass_* had a higher growth rates at the hot edge and better survival and lifespan at the cold edge.

We found limited relationships between wood density, leaf size or xylem vulnerability to embolism and demographic responses at the climatic edge, which was surprising as the mechanisms related to climate response are better understood for these traits. Smaller leaves were related to a longer lifespan and a better survival at the hot edge and a better passage time at the cold edge. This in agreement with Wright et al. (2017) who proposed that large leaves are disadvantaged in hot and dry climates because their transpiration rate during the day is too high and are disadvantaged in cold climate because they have greater risks of reaching critical low temperatures during the night. Anderegg et al. (2019) also reported weak links between traits and drought-related mortality at the edge, with only an effect for xylem vulnerability to embolism. The effect was, however, that drought-adapted species experienced higher drought mortality at the edge (Anderegg et al., 2019). In this study we found no link between xylem vulnerability to embolism and survival response at the edge. In contrast a low xylem vulnerability to embolism (drought-adapted species) was related to better growth at the hot edge (Supplementary Information, Fig. 22).

Finally, our traits analysis might underestimate the role of traits because we ignore intraspecific traits variability. Traits phenotypic plasticity and local adaptation might however be large for species with a broad distribution (see for instance results for *Pinus sylvestris* in Reich, Oleksyn, & Tjoelker, 1996).

### On the challenge of connecting population dynamics and species ranges

A key limitation of our analysis is that it did not include the regeneration phase, which is considered a bottleneck in tree population dynamics and is key to cover the full life cycle to estimate population growth rate (Grubb, 1977). Thus we can not conclude whether our estimates of adult growth and survival are crucial drivers of the population growth rate. In the Supporting Information, we provide an evaluation of the relative importance of the regeneration phase for tree population growth rate with an elasticity analysis of matrix population models extracted from the COMPADRE Plant matrix database (Salguero-Gómez et al., 2015). The elasticity analysis showed that the regeneration and adult phases were equally important (see Fig. 25 in Supporting Information). Our IPMs analysis thus captures an important part of a tree’s life cycle for the population growth rate. However, we can not rule out the possibility that the regeneration phase has a disproportional importance for the dynamics at the edge, as several studies have shown that this phase is extremely sensitive to climate (Canham & Murphy, 2016; Clark, Bell, Kwit, & Zhu, 2014; Defossez, Courbaud, Lasbouygues, Schiffers, & Kunstler, 2016). Integrating fecundity and juvenile lifestages in tree-IPMs is challenging because we have much less data on them (Needham, Merow, Chang-Yang, Caswell, & McMahon, 2018; Ruiz-Benito et al., 2020; but see Lines, Zavala, Ruiz-Benito, & Coomes, 2019).

It is also important to keep in mind that species ranges are not necessarily only related to the mean population growth rate but could also be related to other processes controlling extinction risk. For instance, the temporal variability of population growth rate and the population resilience to disturbances could be crucial at the edge (Holt et al., 2005) but it was not possible to evaluate these processes in our study with the NFI data. Another explanation is that suitable habitats where population growth rates are unaffected might exist up to the edge due to the presence of suitable microsites (Cavin & Jump, 2017). In this case, the species edges arise because the fraction of suitable habitats available to the metapopulation decreases (Holt & Keitt, 2000).

Finally, tree species distributions might not be in equilibrium with the current climate. This could be because species are either still in the process of recolonising from their ice age refugia (Svenning & Skov, 2004) or already affected by climate change. Such disequilibrium should however be visible by better performance at the cold edge (Talluto, Boulangeat, Vissault, Thuiller, & Gravel, 2017) and we found no evidence for this in our results.

### Synthesis

Our study shows that trees’ demographic responses at range edges are more complex than predicted by the ACH. Here, the patterns of demographic response of the 27 European tree species differed between their hot and cold edges. We only found strong evidence of demographic limits for edges occurring in extreme conditions (hot edges of hot-distributed species and cold edges of cold-distributed species). Our findings open an important perspective, as they show that one should not expect the same demographic response at the hot *vs.* the cold edge and that we need to refine predictions of climate change impacts as a function of the edge and species characteristics.

## Acknowledgments

This paper is a joint effort of the working group sAPROPOS - ‘Analysis of PROjections of POpulationS’, kindly supported by sDiv (Synthesis Centre of the German Centre for Integrative Biodiversity Research - iDiv), funded by the German Research Foundation (FZT 118). GK and AG received support from the REFORCE - EU FP7 ERA-NET Sumforest 2016 through the call “Sustainable forests for the society of the future”, with the ANR as national funding agency (grant ANR-16-SUMF-0002). N.R. was funded by a research grant from DFG (RU 1536/3-1). N.R. and C.W. acknowledge the support of the German Centre for Integrative Biodiversity Research (iDiv) funded by Deutsche Forschungsgemeinschaft DFG (FZT 118). MAZ and PRB were supported by grant RTI2018-096884-B-C32 (MICINN, Spain). The NFI data synthesis was conducted within the FunDivEUROPE project funded by the European Union’s Seventh Programme (FP7/2007– 2013) under grant agreement No. 265171. We thank Gerald Kandler (Forest Research Institute Baden-Wurttemberg) for his help to format the German data. We thank the MAGRAMA, the Johann Heinrich von Thunen-Institut, the Natural Resources Institute Finland (LUKE), the Swedish University of Agricultural Sciences, and the French Forest Inventory (IGN) for making NFI data available. We are grateful to the Glopnet, the global wood density, the global leaf size, and the global xylem embolism vulnerability data bases for making their data publicly available. We are grateful to all the participants of the sAPROPOS working group for their stimulating discussion. We are grateful to Fabian Roger for his help to build the species phylogeny.

