## Supporting Information for "Demographic performance of European tree species at their hot and cold climatic edges"

**References** **53**

### Introduction

In this document we present the different steps to develop climate dependent IPMs for trees in Europe, the detailed results, and the elasticity analysis of tree population growth to fecundity from the COMPADRE plant matrix database. First, we present the FunDivEUROPE and climatic data. Secondly, we present the method used to fit tree growth and survival models to these data. Thirdly, we present the approach used to build the IPM kernel from these functions. Then, we present the methods used to derive the demographic metrics from the IPM, and present the sensitivity analysis of the prediction of the IPM. Then, we present the results of the IPM at the edge. Finally, we present the COMPADRE elasticity analysis.

### FUNDIV data presentation

In the framework of the FunDivEUROPE FP7 project, national forest inventory data from several countries were harmonised. This data covers 131,043 plots from the National Forest Inventories (NFIs) of Spain (49,268 plots), France (35,967), Germany (31,477), Sweden (11,746) and Finland (2,585). NFIs record information on individual trees in the plot, including species identity, diameter at breast height (dbh), and status (alive, dead, harvested, or ingrowth - *i.e.* recruitment of tree). Survey and plot design varies between countries, but generally includes circular plots with different sampling radii, depending on the size range of the tree (see below). The minimum dbh of trees included in the FunDivEUROPE dataset is 10 cm, which allowed the same threshold in all countries. The plot coordinates are blurred at 1km because of legal restriction.

*Country specific protocols*

*Spanish National Forest Inventory*

We used information from the second and third Spanish NFI (surveyed in the periods 1986-1996 and 1997-2007, respectively). The Spanish NFI plots are located on a 1 km2 grid in forested regions (Villaescusa & Diaz, 1998; Villanueva, 2004). Spanish NFI plots were sampled using a variable radius technique with four concentric circular subplots of radius 5, 10, 15 and 25 m. Within each subplot, trees were included in the sample according to their diameter at breast height (d.b.h.), with trees smaller than 12.4 cm measured in the 5 m radius subplot, those of 12.5-22.4 cm in the 10 m radius subplot, those of 22.5-42.4 cm in the 15 m radius subplot, and those with d.b.h. larger or equal to 42.5 cm in the 25 m radius subplot.

*French National Forest Inventory*

The French NFI is based on a systematic 1 km2 square grid covering the entire country (https://inventaire-forestier.ign.fr/). The whole grid is measured in 5 years and the 5-year sample is divided into five systematic annual subsamples. Trees are measured in three concentric plots, depending on their circumference at 1.3 m. Trees more than 23.5 cm circumference are measured on a 6 m radius plot; trees more than 70.5 cm circumference are measured on a 9 m radius plot; trees more than 117.5cm circumference are measured on a 15 m radius plot; and trees less than 23.5 cm circumference are not measured. For living tree, radial growth over the last five years is measured on short cores.

*German National Forest Inventory*

We used information from the first and second German NFI. The German NFI uses a systematic grid of clusters, sampled in the periods 1986-1990 (undertaken in West Germany only) and 2001-2002. The size of the sample grid is 4 by 4 km, however, it is reduced in some federal states to either 2.83 by 2.83 km or 2 by 2 km. Each cluster is a quadrangle of 150 m in length with a sample plot on each corner (Kändler, 2006). Trees with a d.b.h. of 10 cm or more in the first inventory and 7 cm in the second were selected by the angle-count method with a basal area factor (BAF) of 4 m2 ha-1 if they are alive or recently dead.

*Finnish National Forest Inventory*

We used data from the eighth NFI (NFI8) of Finland sampled in the period 1985-1986 to 1995. The sample plots are in a systematic grid across the country of plot clusters in forested areas (Mäkipää & Heikkinen, 2003). In Southern Finland the grid is 16 by 16 square km, with four plots in each cluster at 400 m intervals, while in Northern Finland the grid is a 24 by 32 km rectangle with three plots per cluster, at 600 m. intervals. These permanent sample plot data were sampled using a variable radius technique with two concentric circular subplots of radius 5.64 m for trees under 10.5 cm of d.b.h. (i.e. 100 m2) and 9.77 m for trees of d.b.h. 10.5 cm or higher (i.e. 300 m2).

*Swedish National Forest Inventory*

The permanent inventory uses a randomly planned regular sampling grid and includes about 4,500 permanent tracts, each surveyed every five years. Plots in the first census were surveyed between 2003 and 2005 and plots in the second census were surveyed between 2008 and 2010. The tracts are rectangular and have different dimensions depending on the location within the country. Each tract has between 4 and 8 circular sample plots. Trees greater than 10 cm d.b.h. are sampled in a 10 m radius.

Table 1: Country specific protocol description.

| Country | Survey dates | Mean time between census | Protocol type | Protocol description |
|---|---|---|---|---|
| Spain | from 1986/1996 to 1997/2007 | 11.1 years | circular plots | concentric circular plots of different radii, based on dbh as follow: radius of 5m for dbh > 7.5 cm, 10m for dbh > 12.5 cm, 15m for dbh > 22.5cm, and 25m for dbh > 42.5cm |
| France | from 2005 to 2011 | 5 years | circular plots | concentric circular plots of different radii, based on dbh as follow: radius of 6m for dbh > 7.5 cm, 9m for dbh > 22.5 cm, and 15m for dbh > 37.5cm |
| Germany | from 1986/1990 to 2001/2002 | 13.7 years | angle count | angle count method with a basal area factor of 4 $m^2 ha^{-1}$ |
| Finland | from 1985/1986 to 1995 | 9.2 years | circular plot | concentric circular plots of different radii, based on dbh as follow: radius of 5.64m for dbh > 0cm, and 9.77m for dbh > 10.5 cm, |
| Sweden | from 2005/2010 to 2008/2010 | 4.4 years | circular plot | one circular plot of radius 10m for dbh > 10cm |

### Climatic data

As described in the main text we focus on two main climatic variables: (1) the sum of growing degree days above 5.5 °C (*sgdd*) and (2) water aridity index (*wai*). The variation of these two variables is presented in the two following maps.

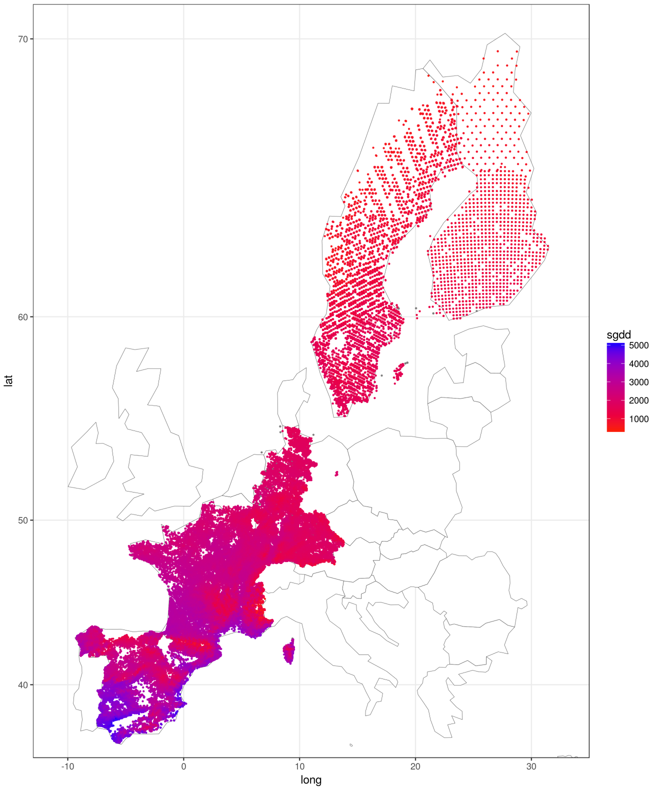

Figure 1: Variation in the sum of growing days (sgdd) across FUNDIV NFI plots.

To describe the species distributions we focused on the axis one of the PCA of these two variables (see Fig. 3). Note that, despite this simplified representation of the range, it was important to have distinct effects of *sgdd* and *wai* in the IPM because they represent different well-known physiological stresses (Kunstler et al., 2011).

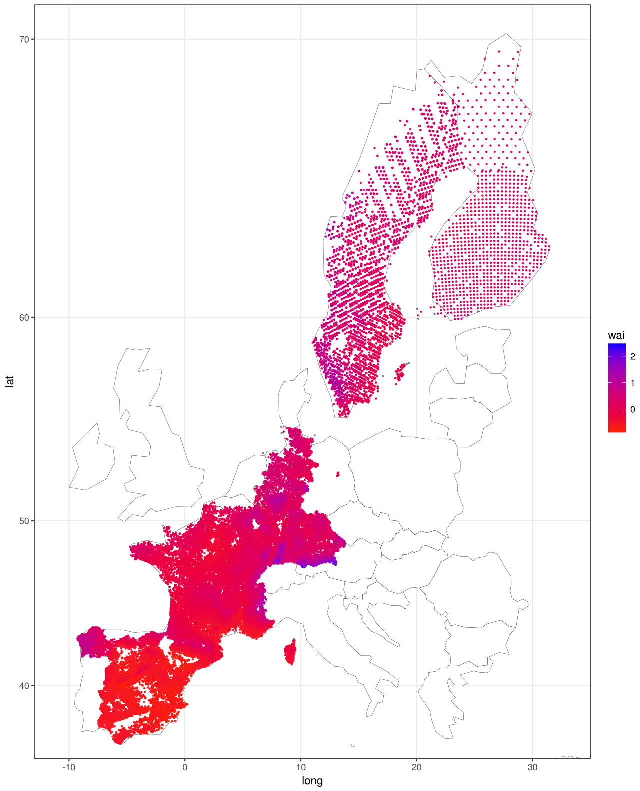

Figure 2: Variation in the water aridity index (wai) across FUNDIV NFI plots.

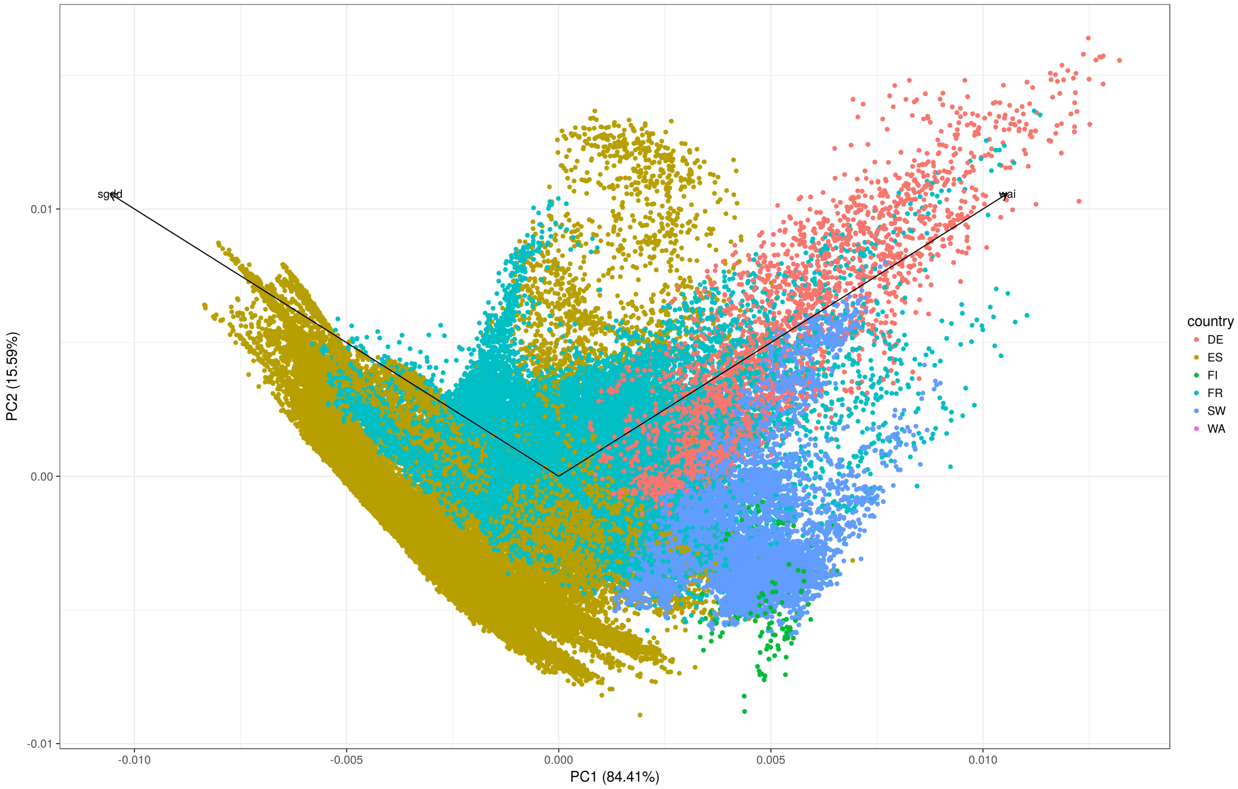

Figure 3: PCA of the two selected climatic variables, sgdd and wai.

### Growth and survival models fitting

The main text presents the methods used to fit the growth and survival models. Here we present the equations of the survival model, the estimated response curves for each co-variable, the re-sampling method and the best model selected for each species.

*Survival model*

The survival models are based on the two following minimal equations:

$$\begin{aligned} cloglog(S_{i,p}) =& a_{0,c} + a_1\, D_i + a_2\, log(D_i) + \\ & a_3\, BA_i + a_4\, \frac{1}{sgdd_p} + a_5\, \frac{1}{wai_p} + y_p \end{aligned} \tag{1}$$

$$\begin{aligned} cloglog(S_{i,p}) =& a_{0,c} + a_1\, D_i + a_2\, log(D_i) + a_3\, BA_i + \\ & a_4\, sgdd_p + a_5\, sgdd_p^2 + a_6\, wai_p + a_7\, wai_p^2 + y_p \end{aligned} \tag{2}$$

Where $S_{i,p}$ is an integer with value 1 if the tree $i$ survived between the two surveys and 0 if the tree died, $y_p$ is the number of years between the two surveys added as an offset. $D_i$ is the diameter at breast height -dbh- of tree $i$, $BA_i$ is the sum of basal area of competitors for tree $i$ per ha, and $sgdd_p$ and $wai_p$ are respectively the sum of growing degree days and the water aridity index of the plot $p$. $a_0$ to $a_7$ are fitted parameters. $a_{0,c}$ is a country specific intercept to account for differences of protocol between countries.

*Data re-sampling*

To account for such uncertainties, we re-sampled 100 times 70% of the data to fit the model and select the best type of response climatic curves and interactions based on the Akaike information criteria (*i.e.*, lowest AIC) (Burnham & Anderson, 2002). Because there were fewer plots in extreme climatic conditions, the estimation maybe biased by disequilibrium of data along climatic gradients with more observation in the core of the climatic niche. Our re-sampling approach re-sampled plots in extreme climatic conditions for the given species with a higher probability: we divided the climatic variables (*sgdd* and *wai*) based on 30 classes of PC1 of the PCA over the range where the species was present and we set the probability of re-sampling for the plots in each class based on the inverse of the number of plots in that class.

*Model selection*

For each 100 resampling of the data, we selected the best growth and survival model between six different competing models (see the percentage of models selected for each species for growth and survival respectively in Table 2 and 3).

*Growth model selection*

For the AIC comparison of the growth models, we first fitted all models based on the maximum likelihood in 'lmer' to select the best model. Then we refitted the best model based on the restricted maximum likelihood because this gives unbiased estimation of variance component.

Table 2: **Growth models selection.** For each of the 6 competing growth model the percentage of selection across the 100 data resampling is indicated. Model names - Asymp: asymptotic response curve, Poly: quadratic polynomial response curve, Inter Clim_Size : interactions between climatic variables and size, Inter Clim_Comp: interactions between climatic variables and competition.

| Species | Asymp | Asymp Inter Clim_Size | Asymp Inter Clim_Size Clim_Comp | Poly | Poly Inter Clim_Size | Poly Inter Clim_Size Clim_Comp |
|---|---|---|---|---|---|---|
| *Abies alba* | 0 | 41 | 0 | 0 | 58 | 1 |

| Species | Asymp | Asymp Inter Clim_Size | Asymp Inter Clim_Size Clim_Comp | Poly | Poly Inter Clim_Size | Poly Inter Clim_Size Clim_Comp |
|---|---|---|---|---|---|---|
| *Acer campestre* | 0 | 0 | 0 | 100 | 0 | 0 |
| *Acer pseudoplatanus* | 0 | 0 | 0 | 72 | 28 | 0 |
| *Alnus glutinosa* | 2 | 37 | 11 | 46 | 4 | 0 |
| *Betula* | 0 | 10 | 10 | 0 | 6 | 74 |
| *Carpinus betulus* | 0 | 100 | 0 | 0 | 0 | 0 |
| *Fagus sylvatica* | 0 | 0 | 0 | 0 | 99 | 1 |
| *Fraxinus excelsior* | 0 | 2 | 46 | 0 | 25 | 27 |
| *Juniperus thurifera* | 0 | 1 | 99 | 0 | 0 | 0 |
| *Larix decidua* | 15 | 0 | 0 | 78 | 7 | 0 |
| *Picea abies* | 0 | 0 | 0 | 0 | 0 | 100 |
| *Pinus halepensis* | 0 | 0 | 0 | 0 | 0 | 100 |
| *Pinus nigra* | 0 | 0 | 0 | 0 | 28 | 72 |
| *Pinus pinaster* | 0 | 3 | 2 | 0 | 0 | 95 |
| *Pinus pinea* | 1 | 60 | 39 | 0 | 0 | 0 |
| *Pinus sylvestris* | 0 | 0 | 100 | 0 | 0 | 0 |
| *Pinus uncinata* | 0 | 0 | 0 | 0 | 5 | 95 |
| *Populus tremula* | 0 | 0 | 0 | 0 | 100 | 0 |
| *Prunus padus* | 1 | 42 | 14 | 9 | 19 | 15 |
| *Quercus faginea* | 0 | 0 | 0 | 47 | 48 | 5 |
| *Quercus ilex* | 0 | 0 | 0 | 0 | 0 | 100 |
| *Quercus petraea* | 0 | 1 | 0 | 0 | 99 | 0 |
| *Quercus pubescens* | 0 | 0 | 0 | 0 | 99 | 1 |
| *Quercus pyrenaica* | 0 | 0 | 0 | 11 | 89 | 0 |
| *Quercus robur* | 0 | 0 | 0 | 0 | 0 | 100 |
| *Quercus suber* | 0 | 0 | 0 | 0 | 1 | 99 |
| *Salix caprea* | 28 | 70 | 0 | 2 | 0 | 0 |

Table 3: **Survival models selection.** For each of the 6 competing survival model the percentage of selection across the 100 data resampling is indicated. Model names as in in Table 2.

| Species | Asymp | Asymp Inter Clim_Size | Asymp Inter Clim_Size Clim_Comp | Poly | Poly Inter Clim_Size | Poly Inter Clim_Size Clim_Comp |
|---|---|---|---|---|---|---|
| *Abies alba* | 94 | 0 | 5 | 1 | 0 | 0 |
| *Acer campestre* | 0 | 0 | 0 | 99 | 0 | 1 |
| *Acer pseudoplatanus* | 92 | 0 | 0 | 8 | 0 | 0 |
| *Alnus glutinosa* | 0 | 0 | 0 | 0 | 100 | 0 |
| *Betula* | 0 | 0 | 0 | 12 | 65 | 23 |
| *Carpinus betulus* | 19 | 1 | 0 | 80 | 0 | 0 |
| *Fagus sylvatica* | 0 | 42 | 0 | 2 | 56 | 0 |
| *Fraxinus excelsior* | 91 | 0 | 8 | 1 | 0 | 0 |
| *Juniperus thurifera* | 72 | 0 | 0 | 25 | 2 | 1 |
| *Larix decidua* | 6 | 47 | 5 | 33 | 0 | 9 |
| *Picea abies* | 0 | 0 | 0 | 0 | 0 | 100 |
| *Pinus halepensis* | 0 | 0 | 0 | 0 | 0 | 100 |
| *Pinus nigra* | 0 | 0 | 0 | 0 | 0 | 100 |
| *Pinus pinaster* | 0 | 0 | 0 | 0 | 0 | 100 |
| *Pinus pinea* | 0 | 95 | 5 | 0 | 0 | 0 |
| *Pinus sylvestris* | 0 | 0 | 0 | 0 | 0 | 100 |
| *Pinus uncinata* | 0 | 0 | 100 | 0 | 0 | 0 |
| *Populus tremula* | 0 | 0 | 0 | 100 | 0 | 0 |
| *Prunus padus* | 13 | 0 | 39 | 0 | 0 | 48 |
| *Quercus faginea* | 0 | 0 | 32 | 0 | 0 | 68 |
| *Quercus ilex* | 0 | 0 | 14 | 0 | 51 | 35 |
| *Quercus petraea* | 100 | 0 | 0 | 0 | 0 | 0 |
| *Quercus pubescens* | 0 | 0 | 0 | 100 | 0 | 0 |
| *Quercus pyrenaica* | 0 | 0 | 9 | 0 | 0 | 91 |

| Species | Asymp | Asymp Inter Clim_Size | Asymp Inter Clim_Size Clim_Comp | Poly | Poly Inter Clim_Size | Poly Inter Clim_Size Clim_Comp |
|---|---|---|---|---|---|---|
| *Quercus robur* | 0 | 0 | 0 | 0 | 0 | 100 |
| *Quercus suber* | 0 | 0 | 0 | 11 | 1 | 88 |
| *Salix caprea* | 32 | 3 | 0 | 65 | 0 | 0 |

*Growth and survival model evaluation*

We evaluated the 100 growth models fitted on a re-sampling of 70% of data by computing the normalised root-mean-square deviation (NRMSD) on the remaining 30% of the data to cross validate the models. The median normalised root-mean-square deviation was below 16 % for most species, except for *Juniperus thurifera* (see Fig. 4), indicating a good predictive power of the growth models.

We evaluated the 100 survival models fitted from re-sampling 70% of data by computing the area under the curve (AUC) on the remaining 30% of the data to cross validate the models. The median AUC ROC, area under the receiver operating characteristic curve, of the survival models was above 0.6 for all species, except for *Salix caprea*, *Quercus suber*, *Pinus pinaster*, and *Pinus halepensis* (see Fig. 5), indicating a fit of medium quality for survival, which is a notoriously difficult process to fit.

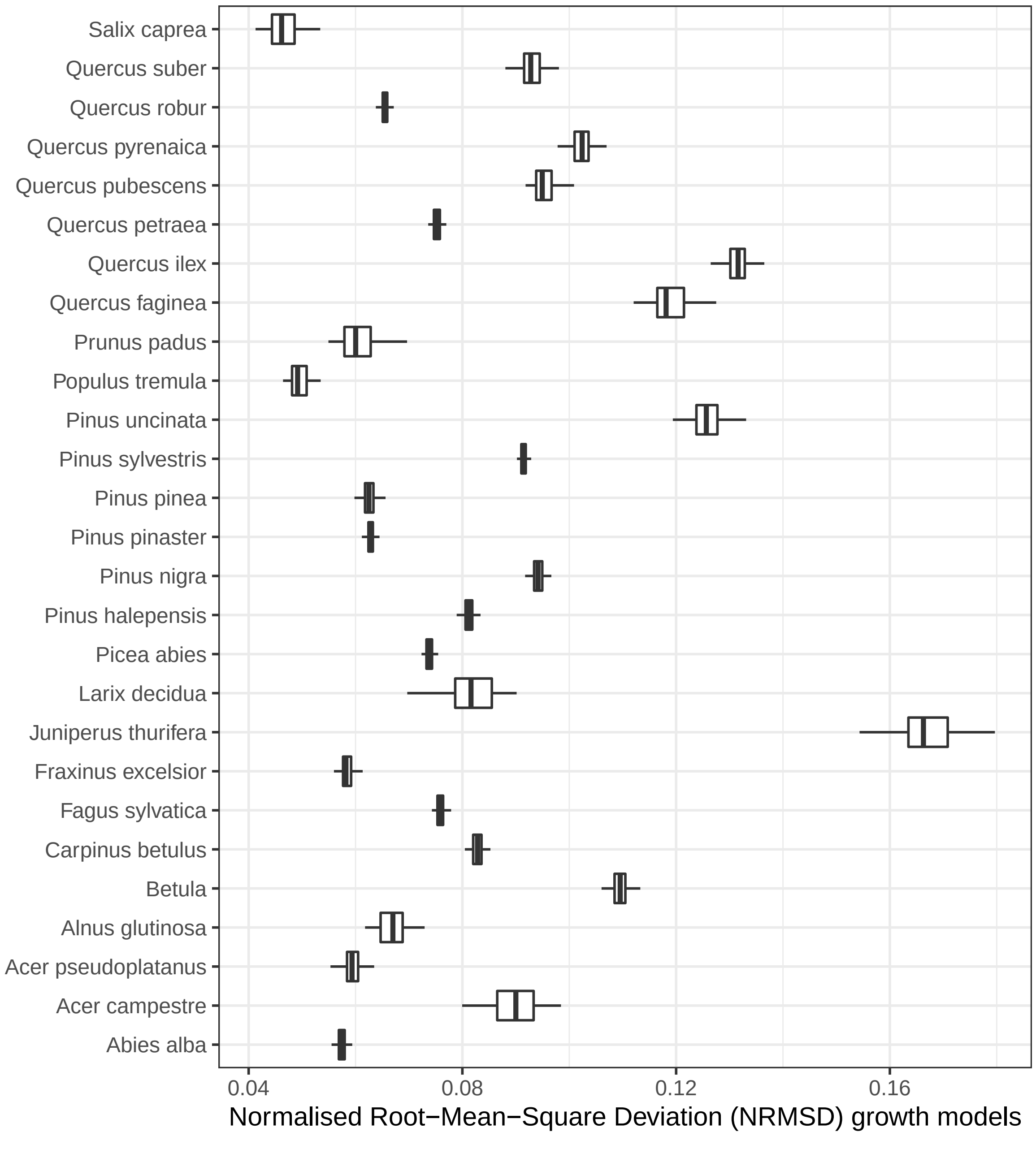

Figure 4: Range of normalised root-mean-square deviation over the 100 resampled growth models per species.

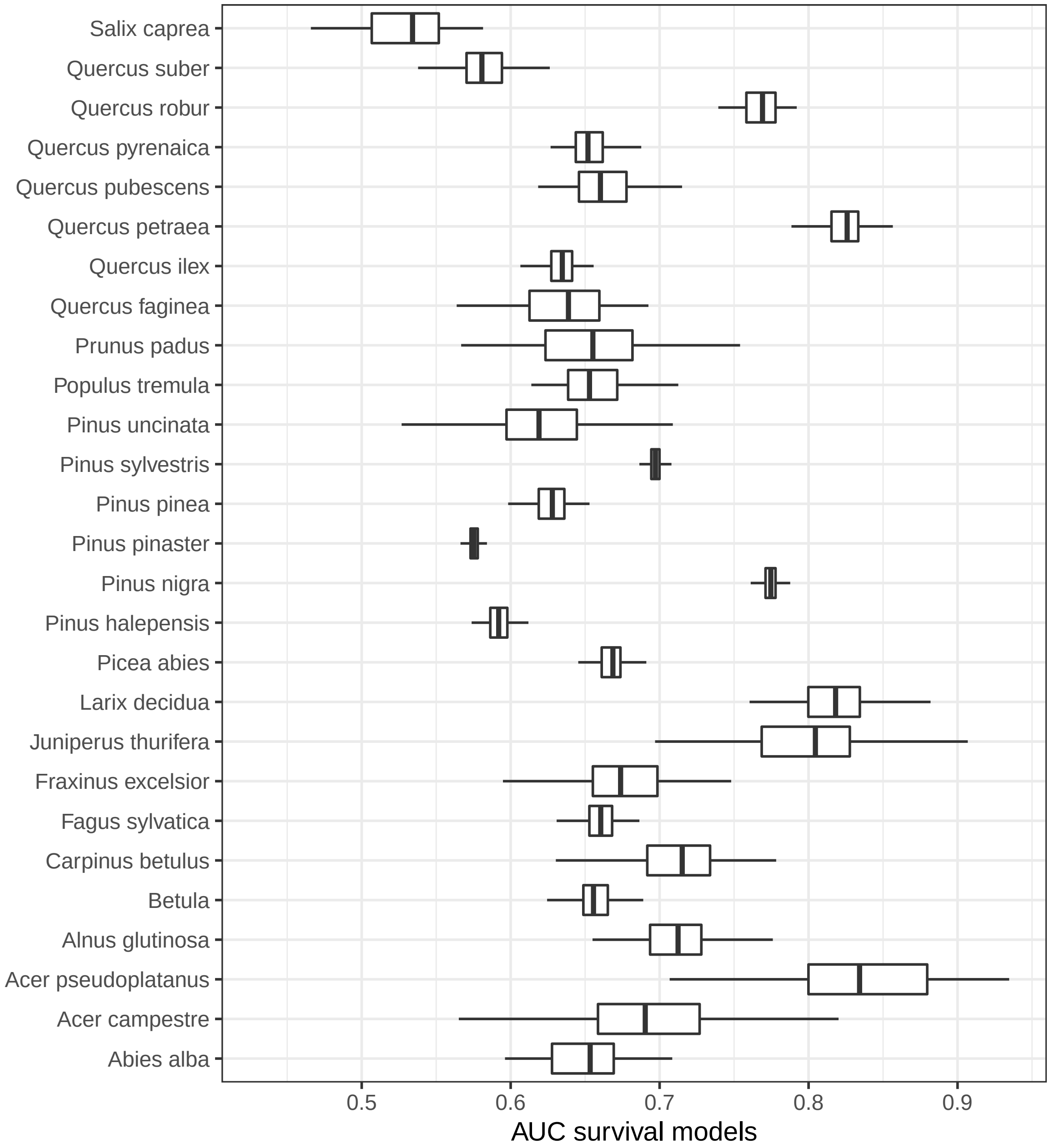

Figure 5: Range of AUC over the 100 resampled survival models per species.

*Variables importance*

For each species we estimated the relative importance of climatic variables ($sgdd$, $wai$, and their respective transformation), competition ($BA$), and size ($D$ and $log(D)$) for both growth and survival models with permutation of the explicative variables with the function 'features_importance' from the R package 'ingredients'. This was done with a single best selected model fitted to the full dataset. We rescaled the importance score to vary between 0 and 1. For both growth and survival, the most important variable is generally tree diameter, followed by competition ($BA$), and then the climatic variables (see Tables 4 and 5).

Table 4: **Variable importance in growth models for each species.** Given is the relative importance of the groups of variables over the overall importance off all variables.

| species | climate | BATOTcomp | size |
|---|---|---|---|
| *Pinus sylvestris* | 0.106 | 0.239 | 0.655 |
| *Pinus pinaster* | 0.237 | 0.262 | 0.5 |
| *Picea abies* | 0.278 | 0.325 | 0.397 |
| *Fagus sylvatica* | 0.224 | 0.232 | 0.544 |
| *Quercus ilex* | 0.286 | 0.292 | 0.422 |
| *Pinus nigra* | 0.182 | 0.367 | 0.451 |
| *Pinus halepensis* | 0.179 | 0.35 | 0.471 |
| *Quercus robur* | 0.109 | 0.355 | 0.536 |
| *Quercus petraea* | 0.102 | 0.215 | 0.683 |
| *Betula* | 0.225 | 0.262 | 0.513 |
| *Quercus pubescens* | 0.11 | 0.33 | 0.561 |
| *Quercus pyrenaica* | 0.164 | 0.339 | 0.497 |
| *Abies alba* | 0.0764 | 0.424 | 0.499 |
| *Carpinus betulus* | 0.0795 | 0.321 | 0.599 |
| *Quercus suber* | 0.273 | 0.33 | 0.397 |
| *Pinus pinea* | 0.0464 | 0.277 | 0.676 |
| *Fraxinus excelsior* | 0.051 | 0.341 | 0.608 |
| *Pinus uncinata* | 0.035 | 0.162 | 0.803 |
| *Quercus faginea* | 0.133 | 0.417 | 0.45 |
| *Alnus glutinosa* | 0.101 | 0.334 | 0.565 |
| *Juniperus thurifera* | 0.137 | 0.256 | 0.607 |
| *Populus tremula* | 0.268 | 0.301 | 0.431 |
| *Acer campestre* | 0.247 | 0.356 | 0.397 |
| *Acer pseudoplatanus* | 0.109 | 0.371 | 0.52 |
| *Larix decidua* | 0.0797 | 0.31 | 0.611 |
| *Salix caprea* | 0.149 | 0.344 | 0.507 |
| *Prunus padus* | 0.101 | 0.416 | 0.483 |

Table 5: **Variables importance in survival models for each species.** Given is the relative importance of the groups of variables over the overall importance off all variables.

| species | climate | BATOTcomp | size |
|---|---:|---:|---:|
| *Pinus sylvestris* | 0.116 | 0.38 | 0.504 |
| *Pinus pinaster* | 0.104 | 0.218 | 0.678 |
| *Picea abies* | 0.147 | 0.33 | 0.524 |
| *Fagus sylvatica* | 0.0505 | 0.204 | 0.745 |
| *Quercus ilex* | 0.106 | 0.368 | 0.526 |
| *Pinus nigra* | 0.118 | 0.146 | 0.736 |
| *Pinus halepensis* | 0.0178 | 0.422 | 0.56 |
| *Quercus robur* | 0.157 | 0.271 | 0.572 |
| *Quercus petraea* | 0.108 | 0.159 | 0.733 |
| *Betula* | 0.209 | 0.266 | 0.524 |
| *Quercus pubescens* | 0.102 | 0.277 | 0.621 |
| *Quercus pyrenaica* | 0.135 | 0.419 | 0.446 |
| *Abies alba* | 0.191 | 0.227 | 0.582 |
| *Carpinus betulus* | 0.0332 | 0.178 | 0.789 |
| *Quercus suber* | 0.126 | 0.312 | 0.563 |
| *Pinus pinea* | 0.105 | 0.339 | 0.556 |
| *Fraxinus excelsior* | 0.0368 | 0.119 | 0.845 |
| *Pinus uncinata* | 0.106 | 0.154 | 0.74 |
| *Quercus faginea* | 0.165 | 0.336 | 0.5 |
| *Alnus glutinosa* | 0.022 | 0.153 | 0.825 |
| *Juniperus thurifera* | 0.164 | 0.316 | 0.52 |
| *Populus tremula* | 0.221 | 0.342 | 0.437 |
| *Acer campestre* | 0.0437 | 0.0625 | 0.894 |
| *Acer pseudoplatanus* | 0.201 | 0.291 | 0.508 |
| *Larix decidua* | 0.118 | 0.14 | 0.743 |
| *Salix caprea* | 0.0683 | 0.148 | 0.784 |
| *Prunus padus* | 0.242 | 0.361 | 0.397 |

*Response curves*

For both tree growth and survival response curves showed a maximum at intermediate size (see Figs 6 and 10). The basal area of local competitors had a negative effect for all species on growth and for most species on survival (see Figs 7 and 11). Tree growth and survival response to *sgdd* and *wai* showed either asymptotically increasing or bell shaped response curves for most species (see Figs 8, 9, 12 and 13). Note that the 95% quantile of the growth prediction represents the uncertainty around the mean response curves not the residual variation.

*Growth model responses curves*

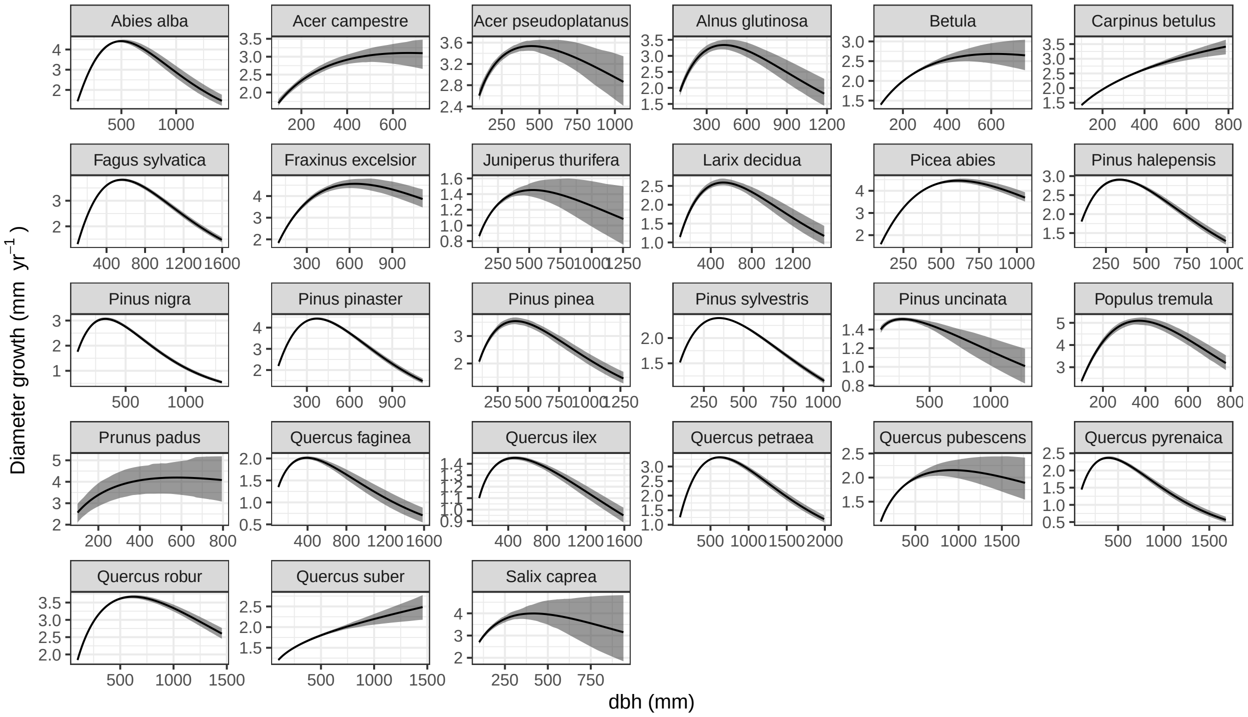

Figure 6: Growth model response curves as a function of dbh per species. The black line represents the mean response over 100 models from resampled data and the grey area the 95% quantile of the growth prediction. Tree dbh varied over the range observed for the species while the other variables were set at their species specific means.

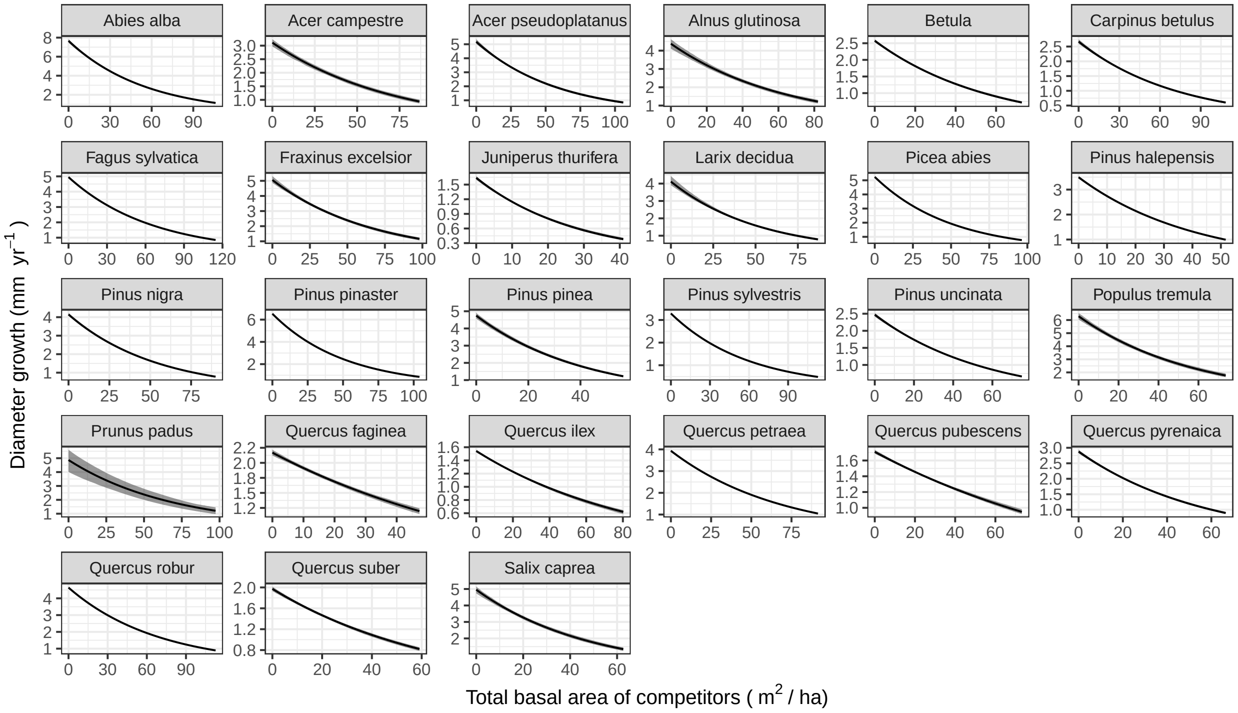

Figure 7: Growth model response curves as a function of total basal area of competitors ($BA$, $m^2 ha^{-1}$) per species. The black line represents the mean response over the 100 models from resampled data and the grey area the 95% quantile of the growth prediction. $BA$ varied over the range observed for the species while the other variables were set at their species specific means.

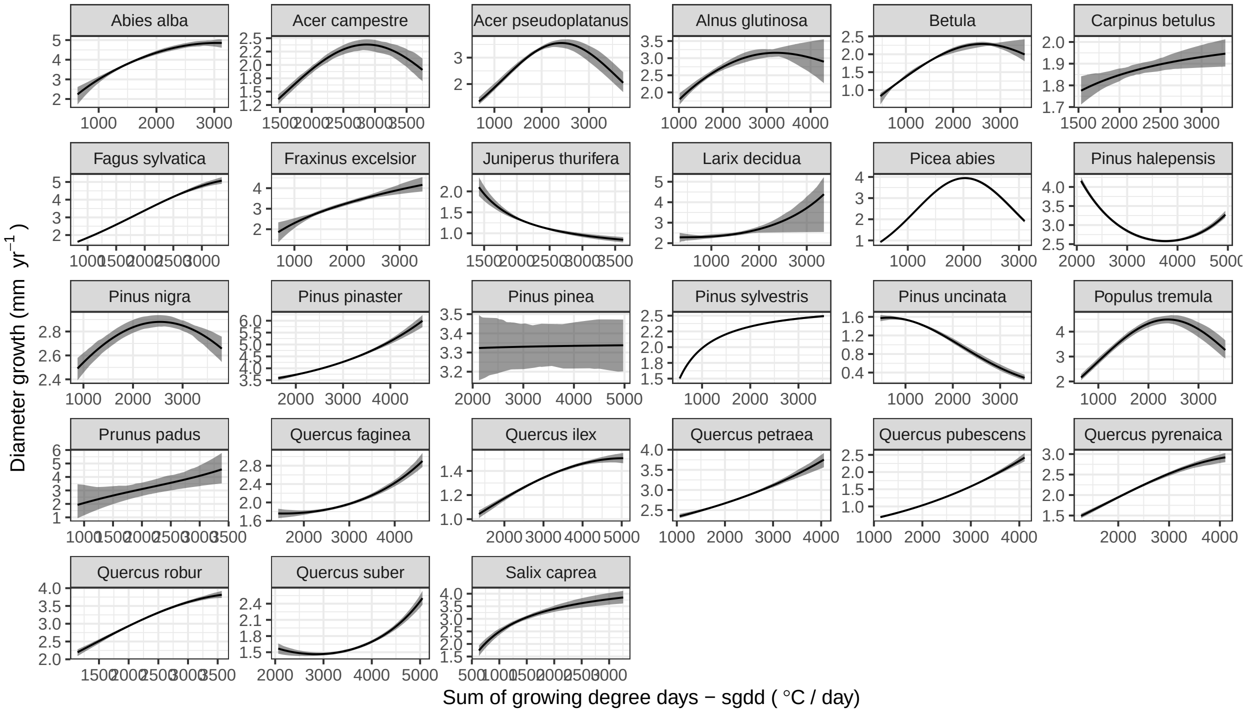

Figure 8: Growth model response curves as a function of sum of growing degree days (*sgdd*, °C) per species. The black line represents the mean response over the 100 models from resampled data and the grey area the 95% quantile of the growth prediction. sgdd varied over the range observed for the species while the other variables were set at their species specific means.

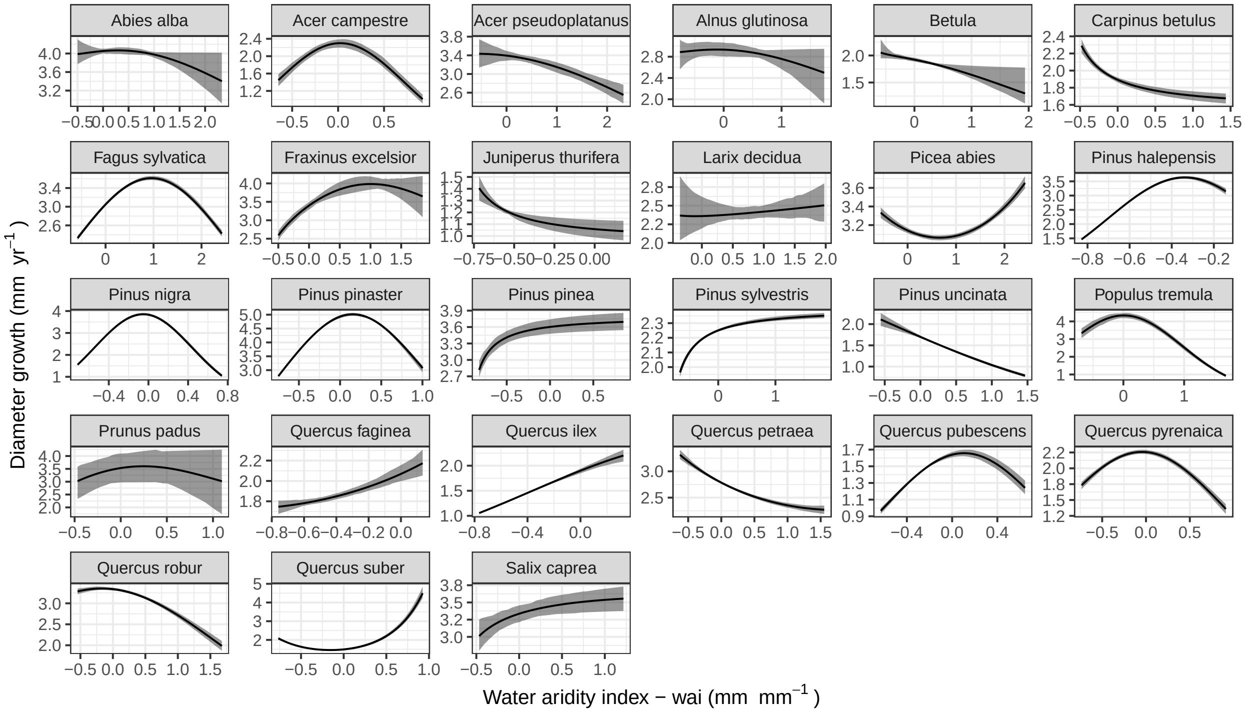

Figure 9: Growth model response curves as a function of the water aridity index (*wai*, mm/mm) per species. The black line represents the mean response over the 100 models from resampled data and the grey area the 95% quantile of the growth prediction. wai varied over the range observed for the species while the other variables were set at their species specific means.

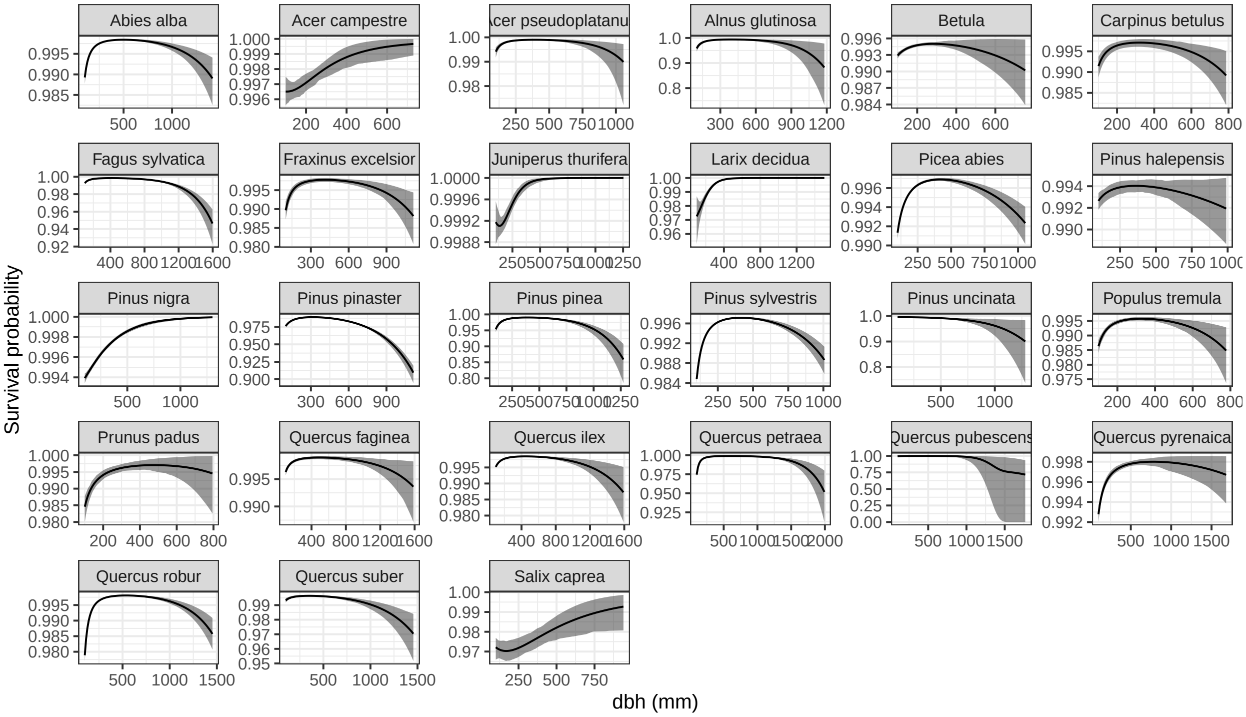

Figure 10: Survival model response curves as a function of dbh per species. The black line represents the mean response over the 100 models from resampled data and the grey area the 95% quantile of the growth prediction. Tree dbh varied over the range observed for the species while the other variables were set at their species specific means.

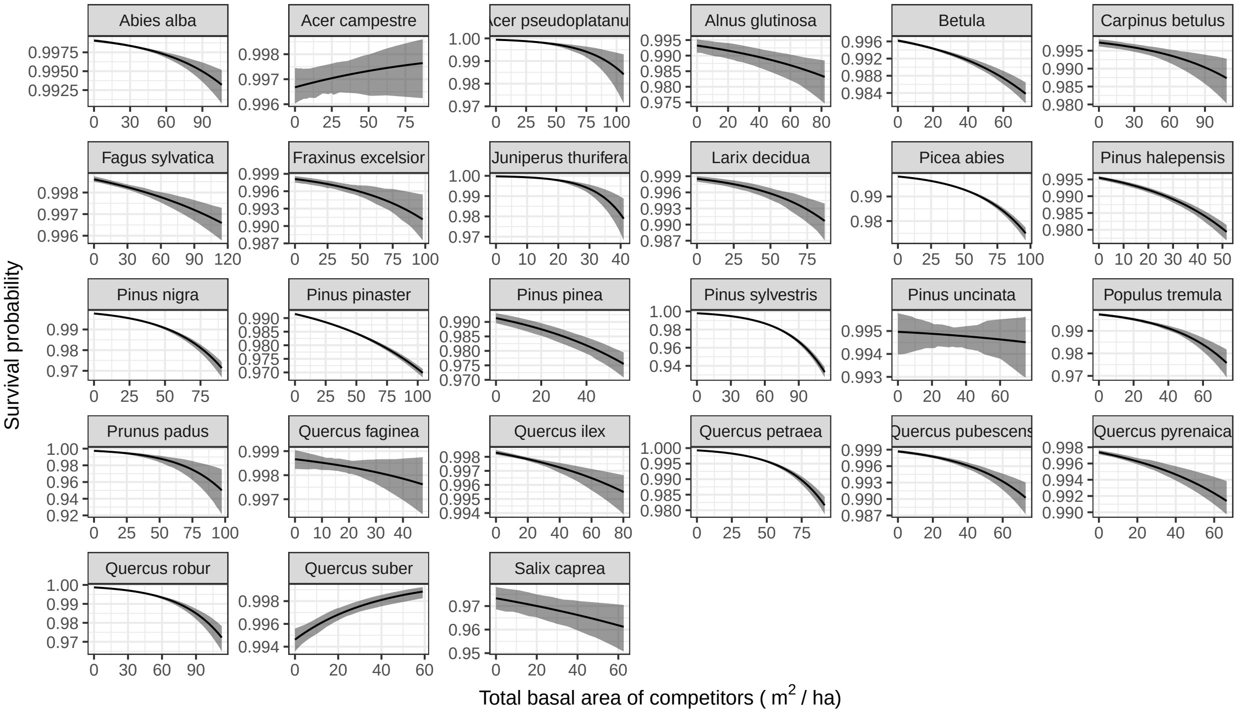

Figure 11: Survival model response curves as a function of total basal area of competitors ($BA$, $m^2ha^{-1}$) per species. The black line represents the mean response over the 100 models from resampled data and the grey area the 95% quantile of the growth prediction. $BA$ varied over the range observed for the species while the other variables were set at their species specific means.

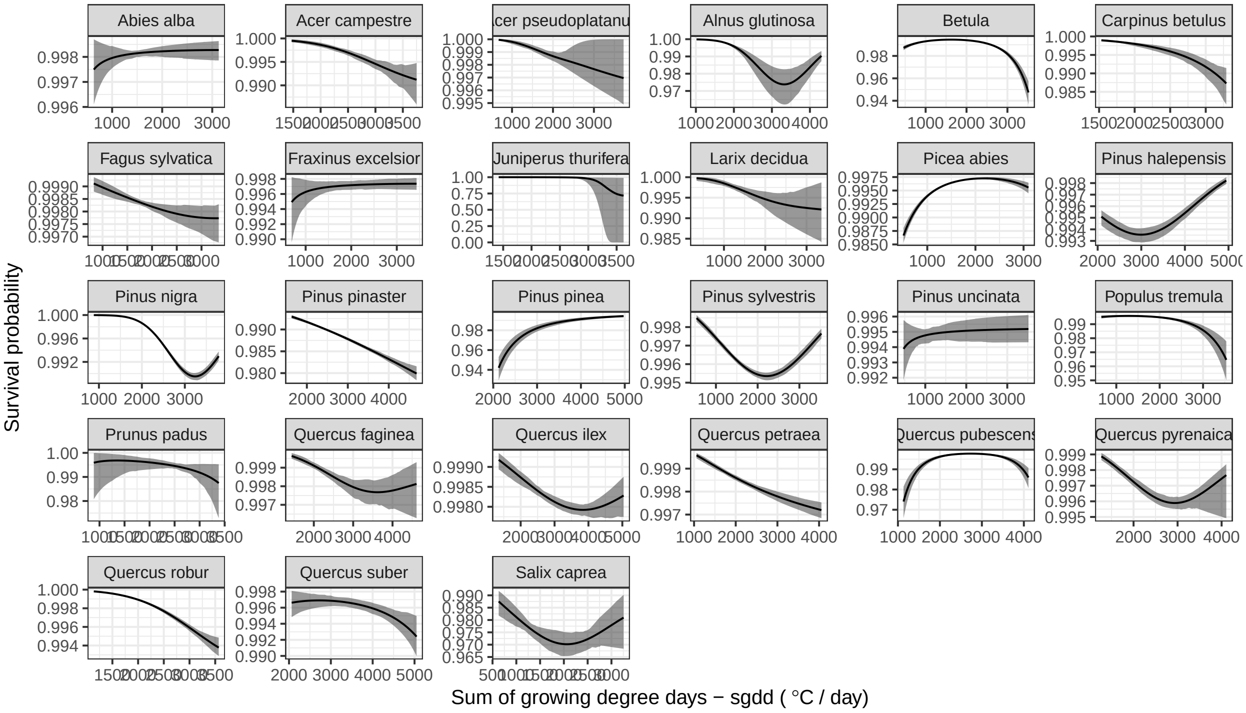

Figure 12: Survival model response curves as a function of sum of growing degree days (*sgdd*, °C) per species. The black line represents the mean response over the 100 models from resampled data and the grey area the 95% quantile of the growth prediction. sgdd varied over the range observed for the species while the other variables were set at their species specific means.

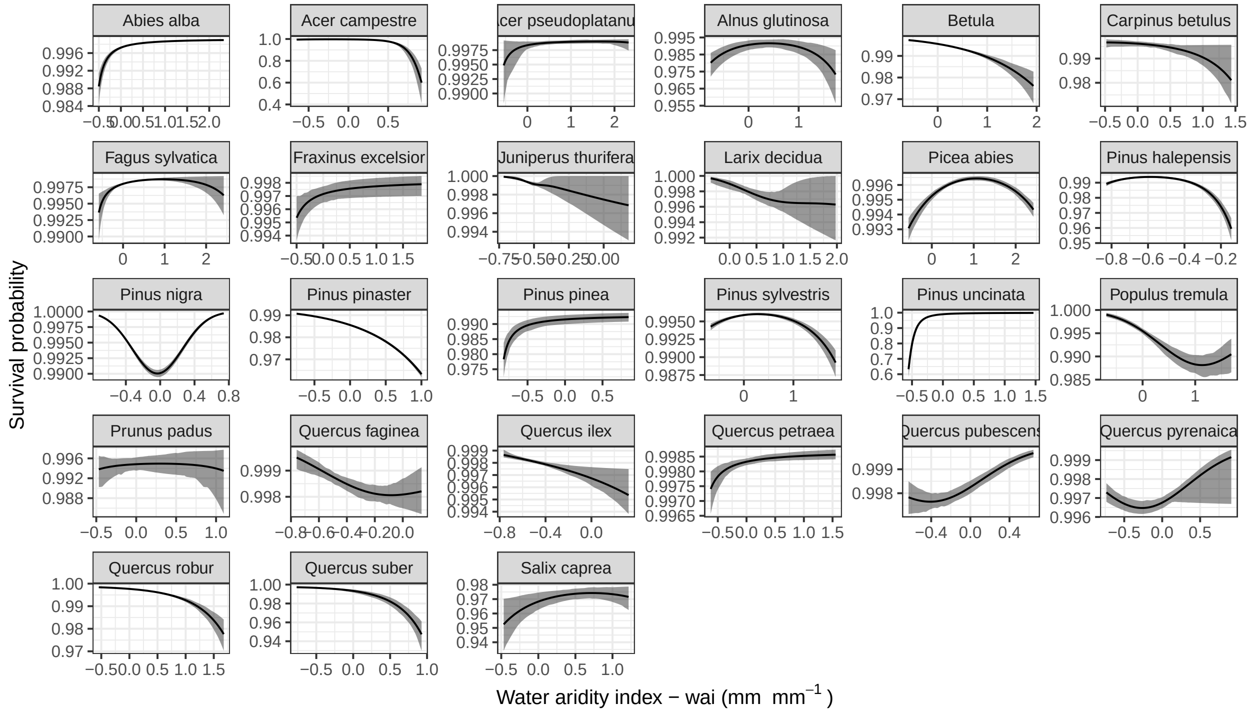

Figure 13: Survival model response curves as a function of the water aridity index (*wai*, mm/mm) per species. The black line represents the mean response over the 100 models from resampled data and the grey area the 95% quantile of the growth prediction. wai varied over the range observed for the species while the other variables were set at their species specific means.

### IPM development

Developing IPMs for trees is notoriously difficult (Needham, Merow, Chang-Yang, Caswell, & McMahon, 2018) because trees are extremely slow growing, resulting in a peaked growth kernel around the diagonal that is difficult to integrate. In order to properly numerically integrate the IPM, while maintaining a reasonable dimension of the IPM matrix and computation time, we mixed mid-bin integration approach and a two dimensions Gauss-Legendre quadrature integration.

#### *Kernel integration*

To develop an effective integration method we combined Gauss Legendre quadrature integration and mid-bin integration. The Gauss Legendre integration was conducted in the vicinity of the diagonal where most of the mass of the growth kernel is distributed and we used a simple mid-bin integration for the reset of the IPM where the growth kernel is flat and close to zero.

We used the 'gauss.quad' function from the R package 'statmod' for the Gauss Legendre integration building on the R code provided in Ellner, Childs, & Rees (2016). Because the shape of the kernel was more complex to integrate in the dimension of size $z'$ at time $t+1$ than on the dimension of the size $z$ at $t$, we only used three nodes for the integration on the dimension of the size at $t$ but 140 nodes for the integration on the dimension at $t+1$. The Gauss Legendre integration was applied till 50 mm from the diagonal.

#### *Change of variable*

In addition, because we fitted the growth model to log diameter growth we needed to recast the growth kernel into a kernel of size at $t+1$ as a function of size at $t$. In order to do this the probability distribution function variable must be changed properly, following (Ellner et al., 2016).

The log diameter growth follow a normal kernel as:

$$f(log(\frac{D_{t+y} - D_t}{y})) \sim \mathcal{N}(\mu, \sigma^2)\,. \tag{3}$$

The kernel of diameter at $t+1$ (annual step), $D_{t+1}$, is then given by:

$$g(D_{t+1}) = f(log(D_{t+1} - D_t)) \times \frac{1}{D_{t+1} - D_t}. \quad (4)$$

### Demographic metrics derivation

Two demographic metrics were derived from the IPM to estimate individual life trajectories that integrate the stochastic processes over the life cycle: (1) the mean lifespan of a 10cm dbh tree and (2) the passage time of a 10cm dbh tree to 60cm (the number of years an individual of 10cm dbh is expected to take to to grow to 60cm dbh).

*Lifespan*

The mean lifespan conditional on initial state was computed following (Ellner et al., 2016):

$$\bar{\eta}(z_0) = \langle e\,(I-P)^{-1}, c \rangle \quad (5)$$

where $e$ is a row matrix of length $m$ the size of the IPM, and $c$ is a vector of initial density of length $m$. Because we computed the lifespan for a tree of 10cm diameter only the first element of $c$ was one and the rest zero.

*Passage time*

The mean passage time to 60cm conditional on initial state was computed as follow (Cochran & Ellner, 1992):

$$\bar{\tau}(z_0) = \frac{(I-P')^{-2}}{(I-P')^{-1}} \quad (6)$$

where $P'$ is the same matrix as $P$ but with zero in the column and row corresponding to the size class 60cm. We extracted the mean passage time for tree of 10cm (the first size class of the IPM)

### IPM diagnostic

To evaluate whether the IPM was leading to eviction (corresponding to individual growing above the maximum size of the IPM being lost) we compared the column sum of the kernel $P$ to the survival function over the size range as they should match without eviction or integration issues. This was the case for all species at extreme values of $sgdd$, $wai$ and $BA$. There was evidence of eviction only in the last upper size classes. This is unavoidable given that our growth kernel can only predict strictly positive growth.

To evaluate if this eviction was problematic we evaluated whether when a population was projected from the minimum size class its abundance decreased to a negligible density before reaching the maximum size class. This was the case for most species and climatic and competitive conditions. This ensured that evication had little effect on our predictions. Finally, we also corrected the IPM for eviction issue using the ceiling eviction correction proposed by Williams, Miller, & Ellner (2012), which in summary consists of adding an additional size class for trees that grow above the maximum size.

*Sensitivity analysis to the IPM size*

To analyse how the demographic metrics were sensitive to the size of the IPM we computed for each species at its median climatic conditions the demographic metrics for IPM of size 50, 100, 200, 500, 700, 800, 1100, 2000, and 3000 (see Fig. 14). We thus chose a size of 800 because the metrics were stable above this size.

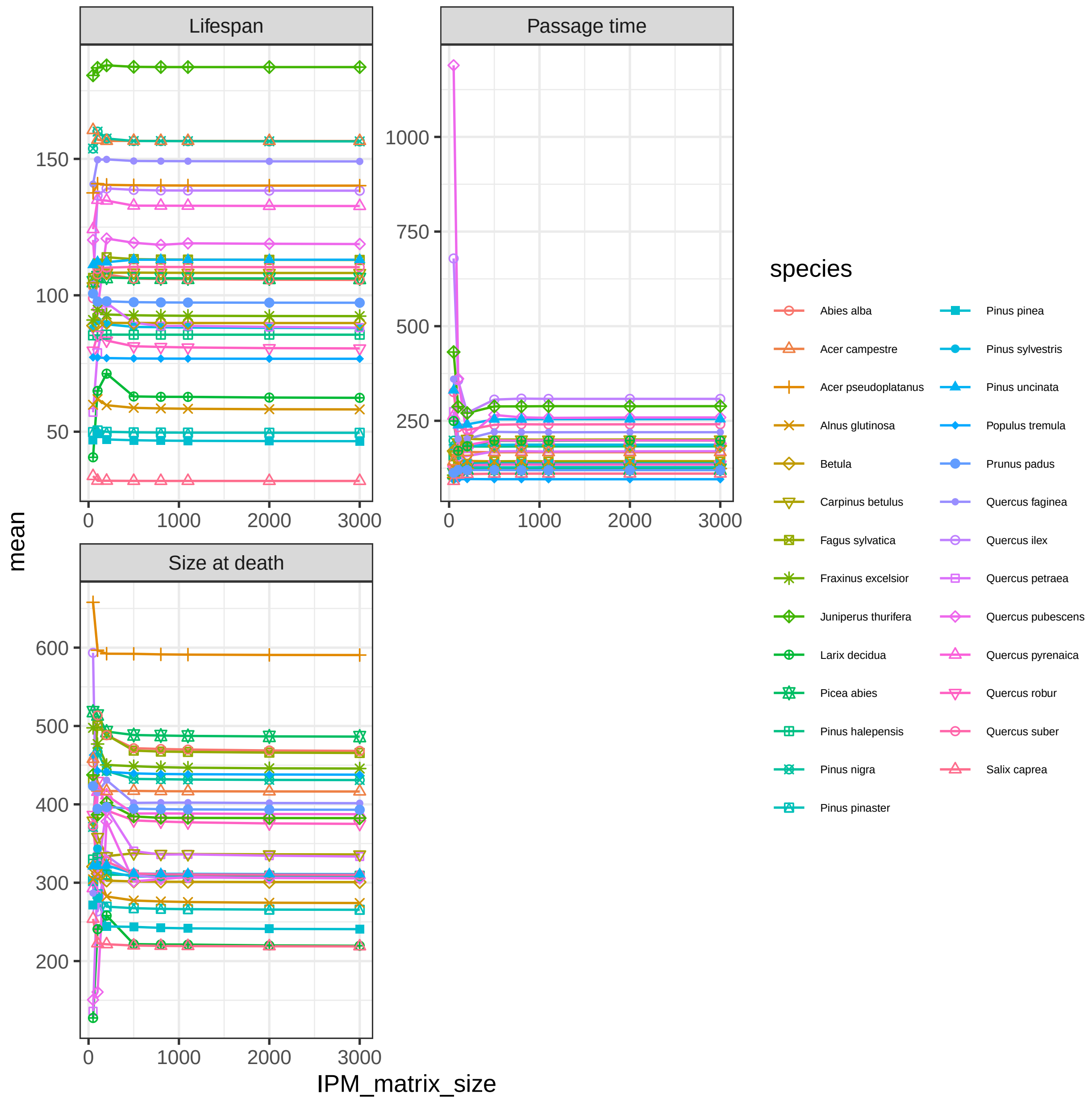

Figure 14: Analysis of the effect IPM size on estimation of lifespan, passage time from 10cm to 60 cm dbh, and size at death.

### Species distribution models

In order to evaluate which species edges in the FunDivEUROPE data correspond to an actual limit of the species distribution and not just a limit in the coverage of the data, we fitted species distribution models with presence/absence data covering all Europe (Mauri, Strona, & San-Miguel-Ayanz, 2017) and retained only the edges for which there was a clear drop in the probability of presence of the species at the edge. We used the EU-Forest data set (Mauri et al., 2017) to fit ensemble species distribution models using four different models in BIOMOD2 (Thuiller, Lafourcade, Engler, & Araújo, 2009). This allow us to have a robust estimation of the probability of presence of each species in each NFI plot. Then, we estimated the mean probability of presence at the center and the two edges by computing the mean probability of presence of the plots that were in climatic bins (as described by the *wai-sgdd* combination) corresponding to the edges or the center. We retained only edges with at least a 10 % drop in the probability of presence at the edge for comparison of the demographic performance at the edge *vs.* the center of the distribution..

*Species distribution data*

Mauri et al. (2017) provides a synthesis of available data on European tree species distribution on a 1 x 1 km grid. A large part of this dataset overlaps with the NFI data used to develop our demographic models, but it also includes NFI data from numerous other countries (including Italy, Norway, Austria, Switzerland, and several eastern European countries) and other data sources, such as the forest focus data base and the Biosoil data base corresponding to a total of 1,000,525 occurrence records.

The FunDivEUROPE NFI dataset only provides a genus-level description for *Betula*, as species-level information are not available in all countries. We thus derived a mean prediction for the two *Betlua* species available in EU-Forest *Betula pendula* and *Betula pubescens*.

*Environmental data*

For each grid point we extracted seven abiotic variables known to influence tree species distribution that were not too strongly correlated. We used the following abiotic variables, mean annual temperature, precipitation of wettest quarter, temperature and precipitation seasonality extracted from the CHELSA (Karger et al., 2017), pH of the first horizon extracted from SoilGrid (Hengl et al., 2017), and an aridity index and actual evapo-transpiration extracted from CGIAR-CSI (Trabucco & Zomer, 2010).

*Ensemble models*

To model species probability of presence we used an ensemble modelling approach based on four different models: generalized linear models with quadratic transformation but no interactions (GLM), general additive models (GAM), generalized boosting model (GBM), and Random Forest. Data was randomly divided into two sets: 70 percent for fitting and 30 percent for evaluation. This was replicated five times per species. We used the True Skill Statistic (TSS) (Hanseen & Kuipers, 1965) metrics to evaluate model performance (Allouche, Tsoar, & Kadmon, 2006). We retained only models in the ensemble model prediction that were above 0.4 TSS. Each individual model weight in the ensemble prediction was proportional to its TSS.

The individual model TSS metrics indicated a high predictive power; only *Salix caprea* and *Prunus padus* had relative poor models (see Fig. 15).

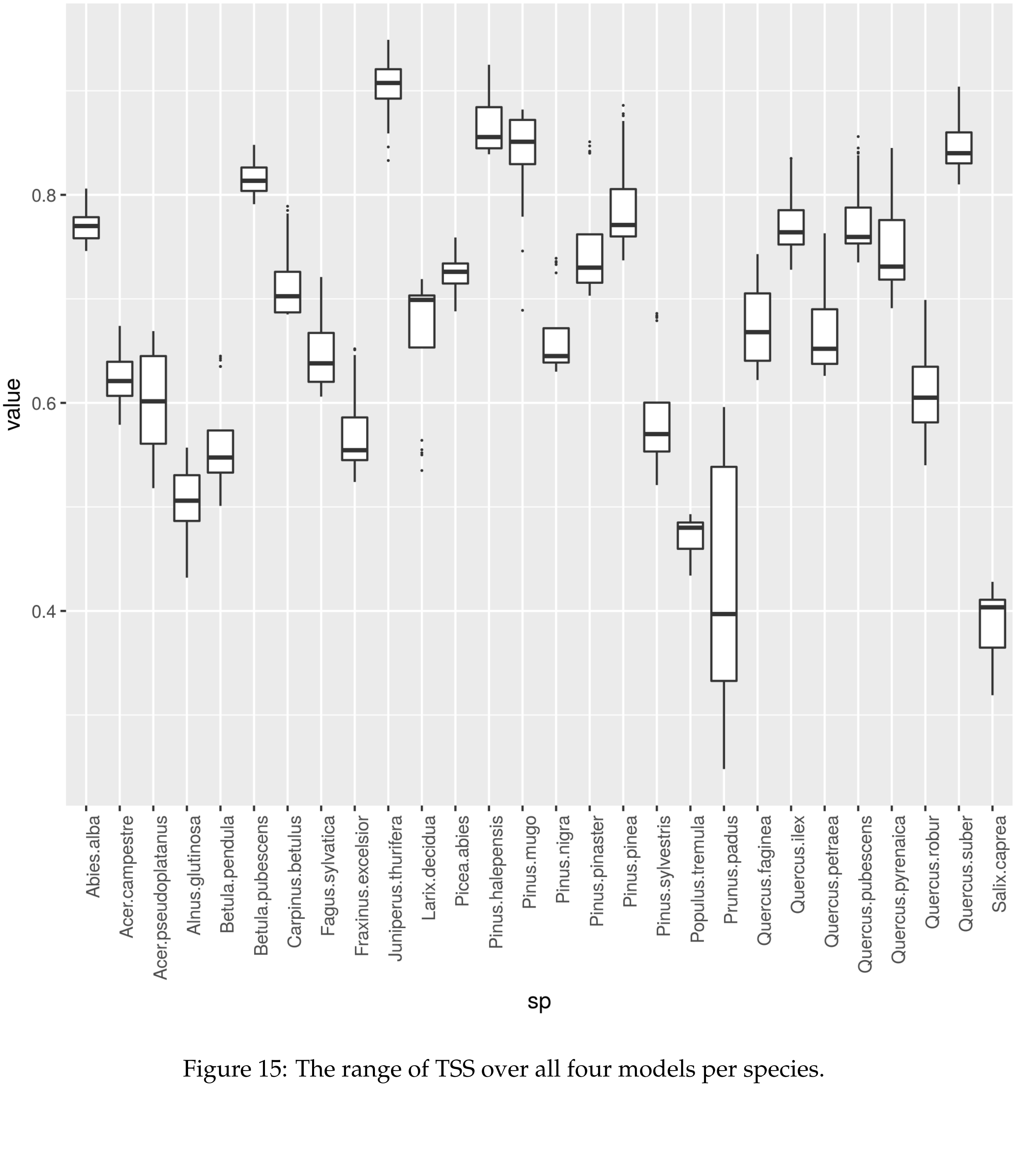

Figure 15: The range of TSS over all four models per species.

### Species demographic performance at the climatic edges

*Species specific demographic metrics at the hot and cold edges* vs. *climatic center*

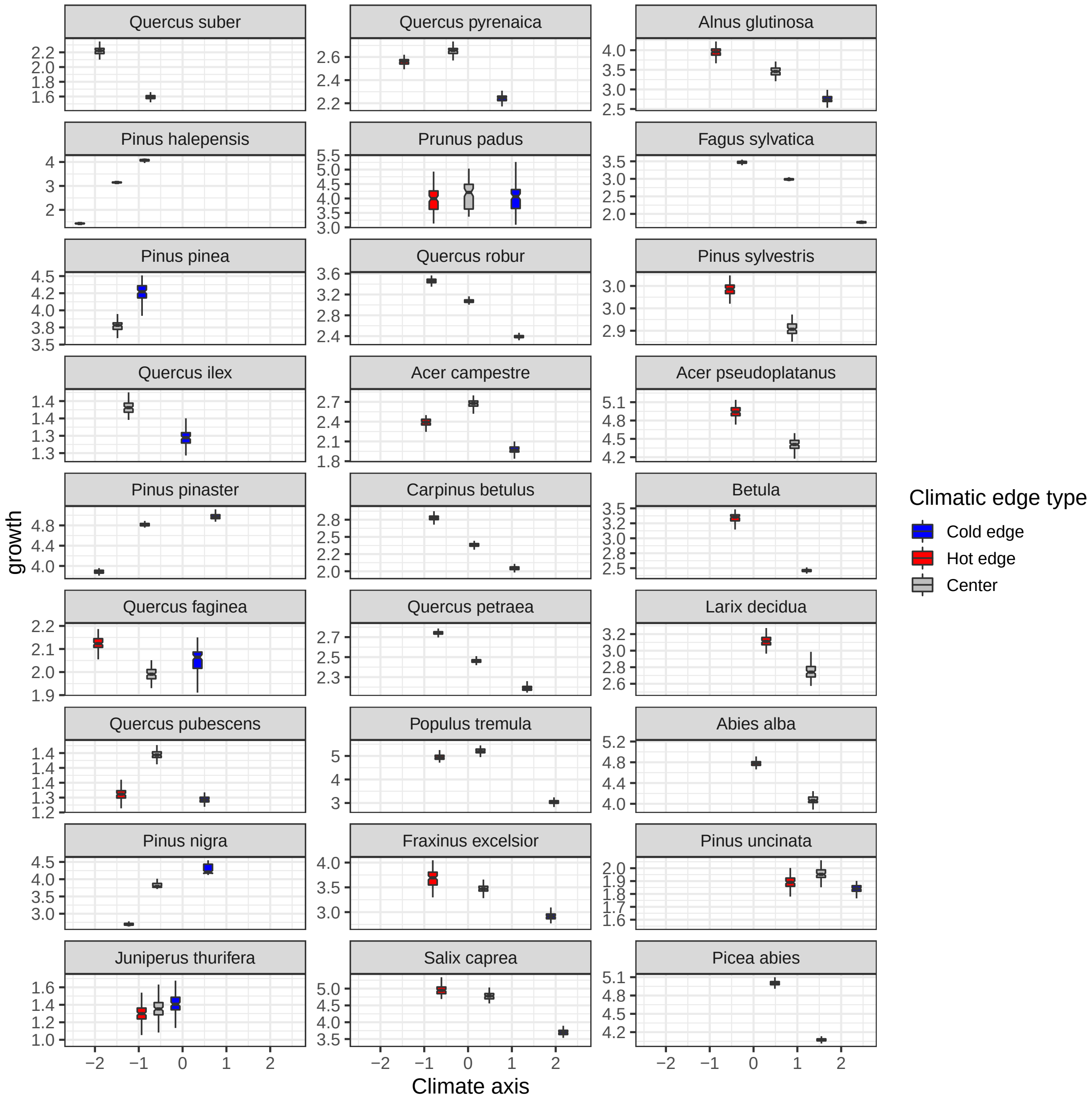

Figure 16: Species demographic metrics at the hot and cold edges and the median climate for the growth of a 15cm dbh tree. Only edges corresponding to a drop in the SDM-derived probability of presence of the species are presented.

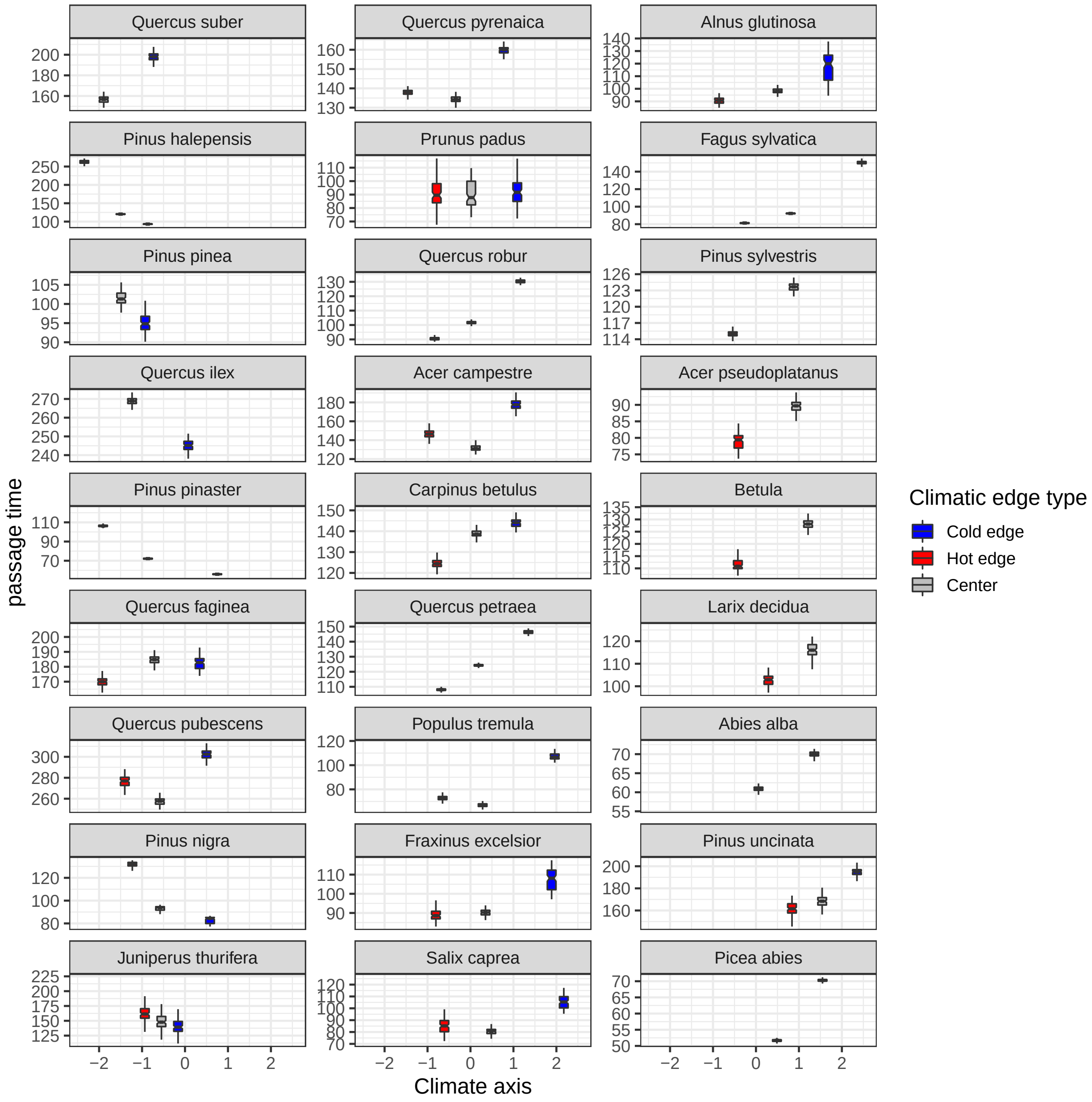

Figure 17: Species demographic metrics at the hot and cold edges and the median climate for the passage time of a 10cm dbh tree to 60cm dbh. Only edges corresponding to a drop in the SDM-derived probability of presence of the species are presented.

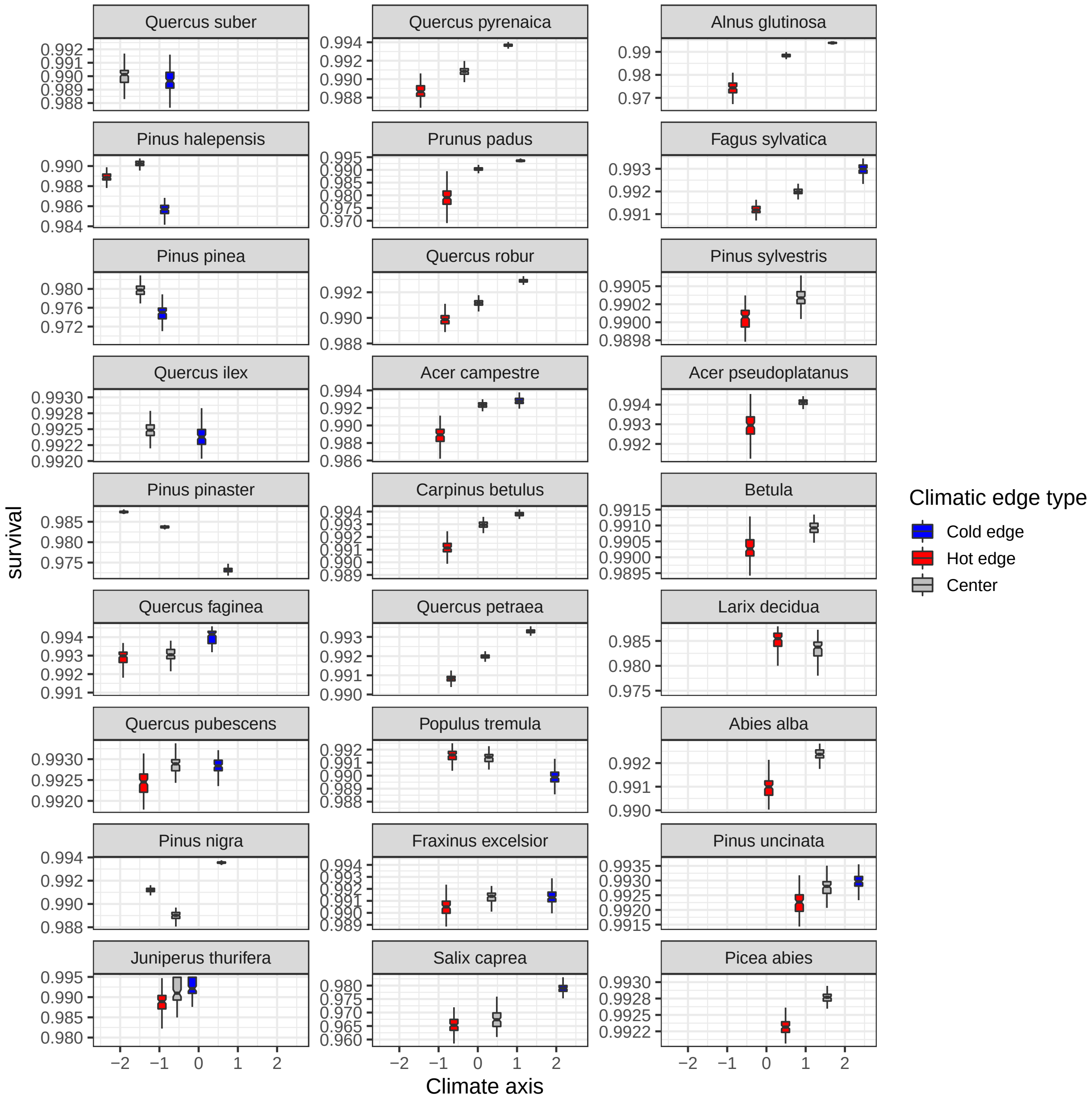

Figure 18: Species demographic metrics at the hot and cold edges and the median climate for the survival rate of a 15cm dbh tree. Only edges corresponding to a drop in the SDM-derived probability of presence of the species are presented.

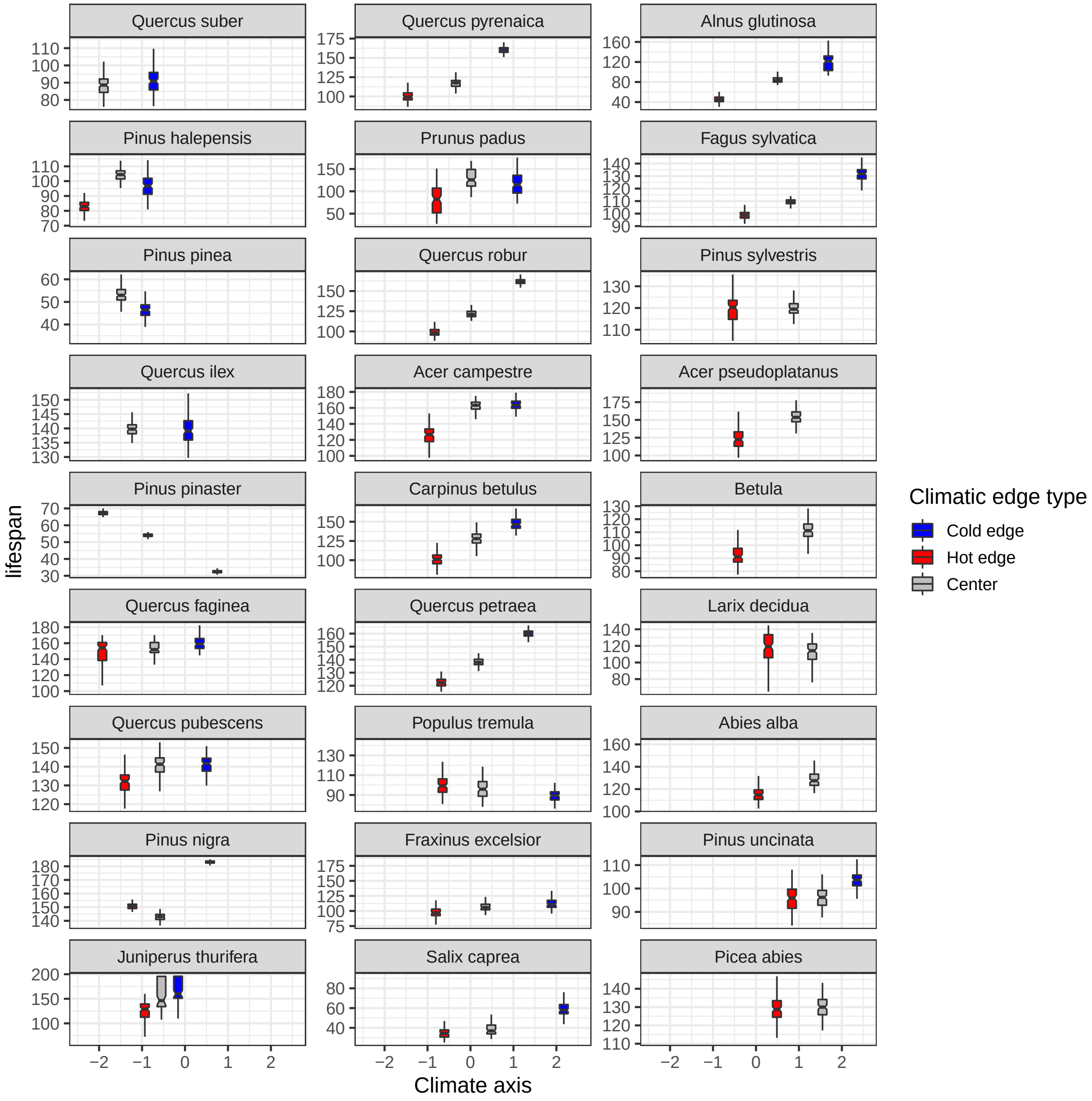

Figure 19: Species demographic metrics at the hot and cold edges and the median climate for the lifespan of a 10cm dbh tree. Only edges corresponding to a drop in the SDM-derived probability of presence of the species are presented.

*Demographic response at the edge in function of species median climate at a high level of competition*

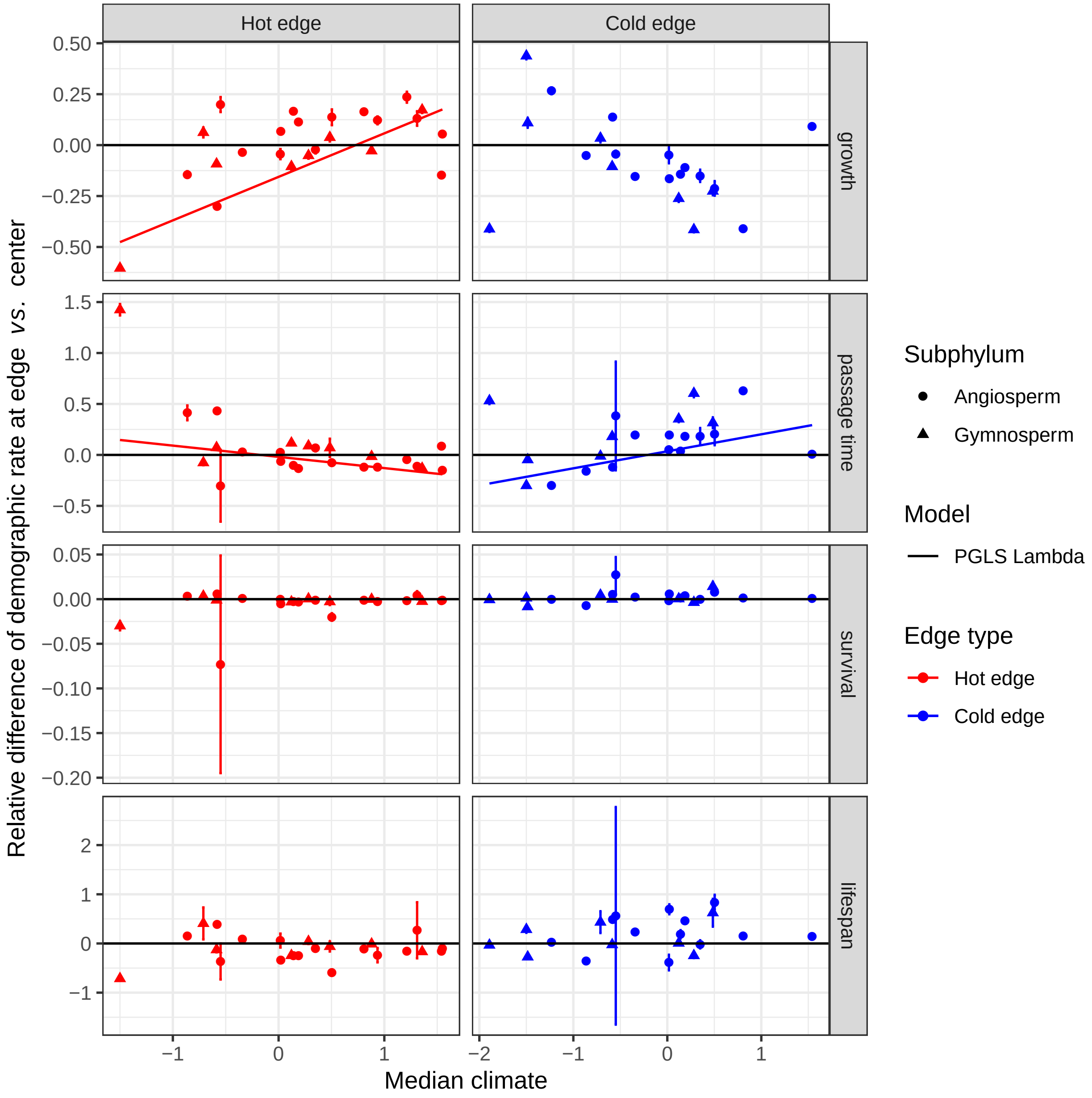

Figure 20: **Changes in demographic responses at the edge ($\Omega^{m}_{edge}$) in function of species median climate with a high level of competition (basal area of competitors = 30 $m^2ha^{-1}$).** Species demographic response at the edge - measured as the relative differences of the demographic metrics at climatic edge *vs.* the median climate of the species distribution - in function of the median position of the species on the first axis of the climate PCA. For each species the mean (point) and the 95% quantiles (error bar) of the demographic response over the 100 data resampling is represented for both the hot (red) and the cold (blue) edges. Phylogenetic generalized least squares (PGLS) regressions are represented only for significant relationship with a non negligible magnitude of the effect. Continuous lines represent model fitted with a PGLS regression where the best value of Pagel's lambda was estimated, and dashed lines represent model fitted with PGLS model with a Brownian model (corresponding to Pagel's lambda set at 1). Gymnosperm and angiosperm species which represent two distinct groups in the phylogeny are represented with different symbols.

Table 6: **Regression between demographic response at the edge and N per mass (%).** P value, effect size (confidence interval in bracket), and adujsted $R^2$ for 'lm' regressions, and p value and effect size (confidence interval in bracket) for phylogentic 'PGLS' regression.

| Demographic metrics | edge type | p value PGLS | effect size PGLS | PGLS model type |
|---|---|---|---|---|
| growth | cold edge | 4e-04 | -0.499 (-0.738 - -0.26) | PGLS Brownian |
| survival | cold edge | 0.0045 | 0.529 (0.186 - 0.872) | PGLS Brownian |
| passage time | cold edge | < 0.0001 | 0.693 (0.606 - 0.78) | PGLS Lambda |
| lifespan | cold edge | < 0.0001 | 1.243 (0.92 - 1.565) | PGLS Brownian |
| growth | hot edge | < 0.0001 | 1.26 (1.26 - 1.26) | PGLS Lambda |
| survival | hot edge | 0.9886 | 0.002 (-0.229 - 0.232) | PGLS Lambda |
| passage time | hot edge | < 0.0001 | -0.494 (-0.618 - -0.369) | PGLS Brownian |
| lifespan | hot edge | < 0.0001 | -0.938 (-1.181 - -0.695) | PGLS Brownian |

Table 7: **Regression between demographic response at the edge and wood density ($g/cm^3$).** P value, effect size (confidence interval in bracket), and adujsted $R^2$ for 'lm' regressions, and p value and effect size (confidence interval in bracket) for phylogentic 'PGLS' regression.

| Demographic metrics | edge type | p value PGLS | effect size PGLS | PGLS model type |
|---|---|---|---|---|
| growth | cold edge | 0.9942 | -0.002 (-0.448 - 0.445) | PGLS Lambda |
| survival | cold edge | 0.8214 | 0.042 (-0.346 - 0.431) | PGLS Lambda |
| passage time | cold edge | 0.2312 | 0.171 (-0.119 - 0.461) | PGLS Lambda |
| lifespan | cold edge | 0.5285 | 0.213 (-0.482 - 0.909) | PGLS Brownian |
| growth | hot edge | 0.066 | -0.116 (-0.241 - 0.008) | PGLS Lambda |
| survival | hot edge | 0.2062 | 0.145 (-0.086 - 0.376) | PGLS Lambda |
| passage time | hot edge | 0.0323 | -0.216 (-0.413 - -0.02) | PGLS Brownian |
| lifespan | hot edge | 8e-04 | -0.29 (-0.445 - -0.136) | PGLS Brownian |

Table 8: **Regression between demographic response at the edge and $\Psi_{50}$ ($MPa$).** P value, effect size (confidence interval in bracket), and adujsted $R^2$ for 'lm' regressions, and p value and effect size (confidence interval in bracket) for phylogentic 'PGLS' regression.

| Demographic metrics | edge type | p value PGLS | effect size PGLS | PGLS model type |
|---|---|---|---|---|
| growth | cold edge | 0.1557 | 0.58 (-0.247 - 1.408) | PGLS Lambda |
| survival | cold edge | 3e-04 | -0.567 (-0.828 - -0.306) | PGLS Lambda |
| passage time | cold edge | 0.7185 | -0.047 (-0.319 - 0.225) | PGLS Lambda |
| lifespan | cold edge | 0.5242 | -0.248 (-1.057 - 0.562) | PGLS Lambda |
| growth | hot edge | 0.0061 | -0.956 (-1.606 - -0.307) | PGLS Lambda |

| Demographic metrics | edge type | p value PGLS | effect size PGLS | PGLS model type |
|---|---|---|---|---|
| survival | hot edge | 0.1259 | -0.06 (-0.139 - 0.019) | PGLS Brownian |
| passage time | hot edge | 0.0067 | 0.32 (0.1 - 0.54) | PGLS Lambda |
| lifespan | hot edge | 0.107 | 0.396 (-0.094 - 0.885) | PGLS Lambda |

Table 9: **Regression between demographic response at the edge and Leaf area ($cm^2$).** P value, effect size (confidence interval in bracket), and adujsted $R^2$ for 'lm' regressions, and p value and effect size (confidence interval in bracket) for phylogentic 'PGLS' regression.

| Demographic metrics | edge type | p value PGLS | effect size PGLS | PGLS model type |
|---|---|---|---|---|
| growth | cold edge | 0.0566 | -0.318 (-0.646 - 0.01) | PGLS Brownian |
| survival | cold edge | 0.002 | 0.495 (0.217 - 0.774) | PGLS Lambda |
| passage time | cold edge | 0.0417 | 0.234 (0.01 - 0.458) | PGLS Brownian |
| lifespan | cold edge | 0.0265 | 0.796 (0.109 - 1.484) | PGLS Brownian |
| growth | hot edge | 0.0126 | 0.465 (0.113 - 0.816) | PGLS Lambda |
| survival | hot edge | 0.1757 | -0.326 (-0.812 - 0.161) | PGLS Brownian |
| passage time | hot edge | 1e-04 | -0.304 (-0.424 - -0.184) | PGLS Lambda |
| lifespan | hot edge | 1e-04 | -0.742 (-1.05 - -0.434) | PGLS Lambda |

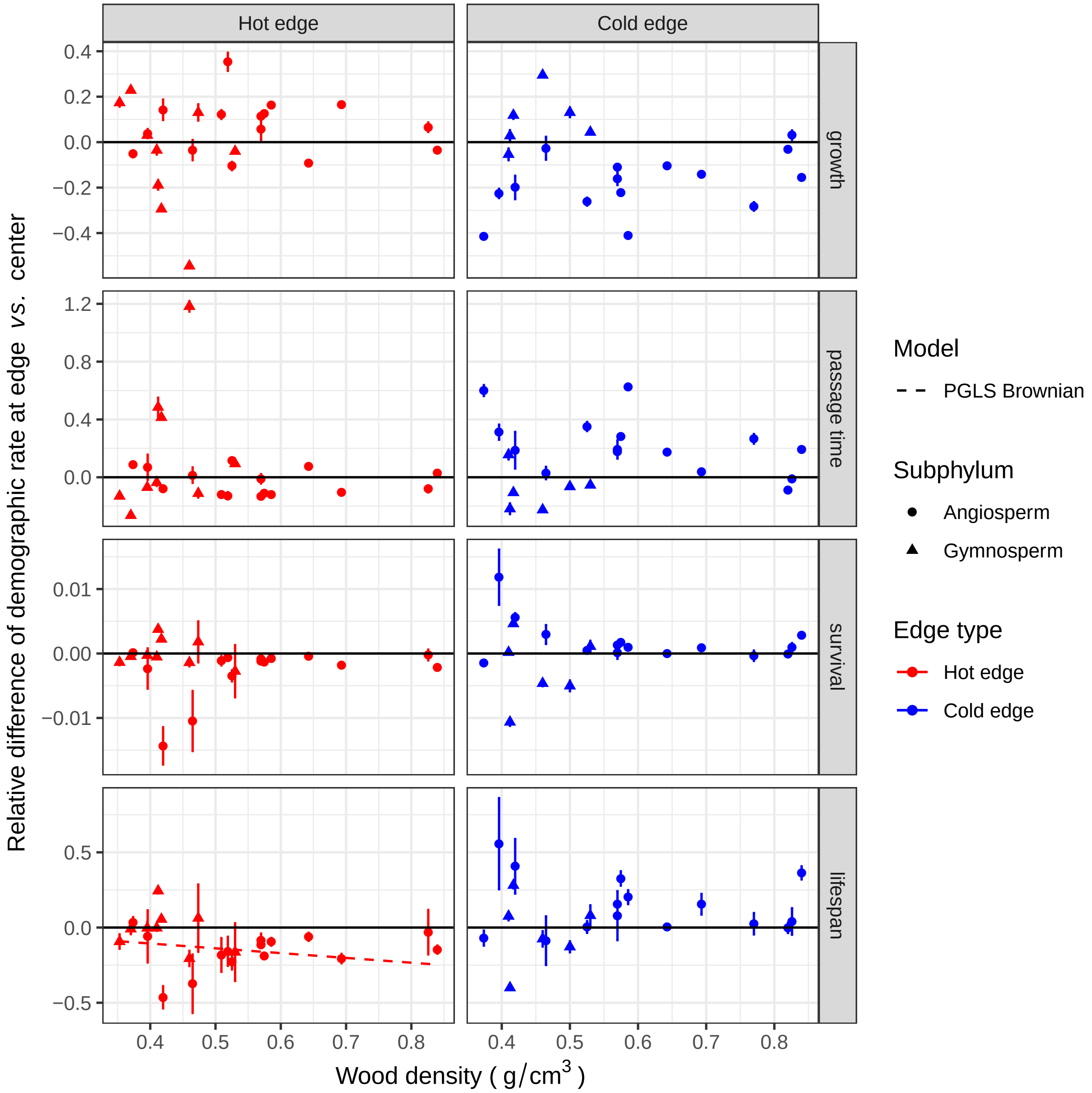

Figure 21: **Changes in demographic responses at the edge ($\Omega^{m}_{edge}$) in function of species wood density.** Species demographic response at the edge - measured as the relative differences in the demographic metric at climatic edge *vs.* the median climate of the species distribution - as a function of species leaf nitrogen per mass. For each species the mean (point) and 95% quantiles (error bar) of the demographic response over the 100 data resampling is represented for both the hot (red) and the cold (blue) edges. Phylogenetic generalized least squares (PGLS) regressions are represented only for significant relationship with a non negligible magnitude of the effect. PGLS regression were first estimated by estimating the best value of Pagel's lambda (a measure of the phylogenetic signal), if the fit failed we fitted a PGLS model with a Brownian model corresponding to Pagel's lambda set at 1.

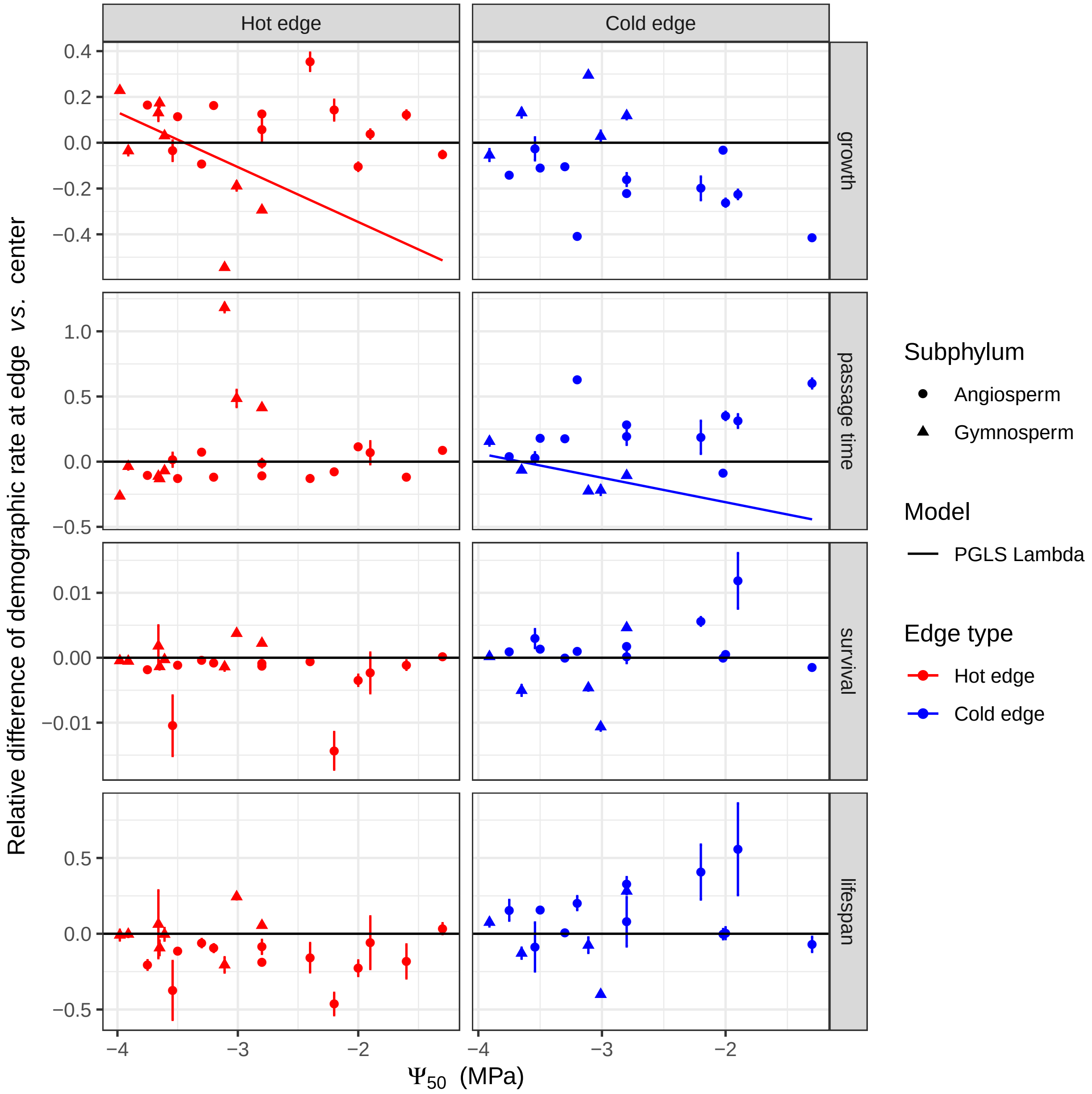

Figure 22: **Changes in demographic responses at the edge ($\Omega^{m}_{edge}$) in function of species $\Psi_{50}$.** Species demographic response at the edge - measured as the relative differences in the demographic metric at climatic edge *vs.* the median climate of the species distribution - as a function of species leaf nitrogen per mass. For each species the mean (point) and 95% quantiles (error bar) of the demographic response over the 100 data resampling is represented for both the hot (red) and the cold (blue) edges. Phylogenetic generalized least squares (PGLS) regressions are represented only for significant relationship with a non negligible magnitude of the effect. PGLS regression were first estimated by estimating the best value of Pagel's lambda (a measure of the phylogenetic signal), if the fit failed we fitted a PGLS model with a Brownian model corresponding to Pagel's lambda set at 1.

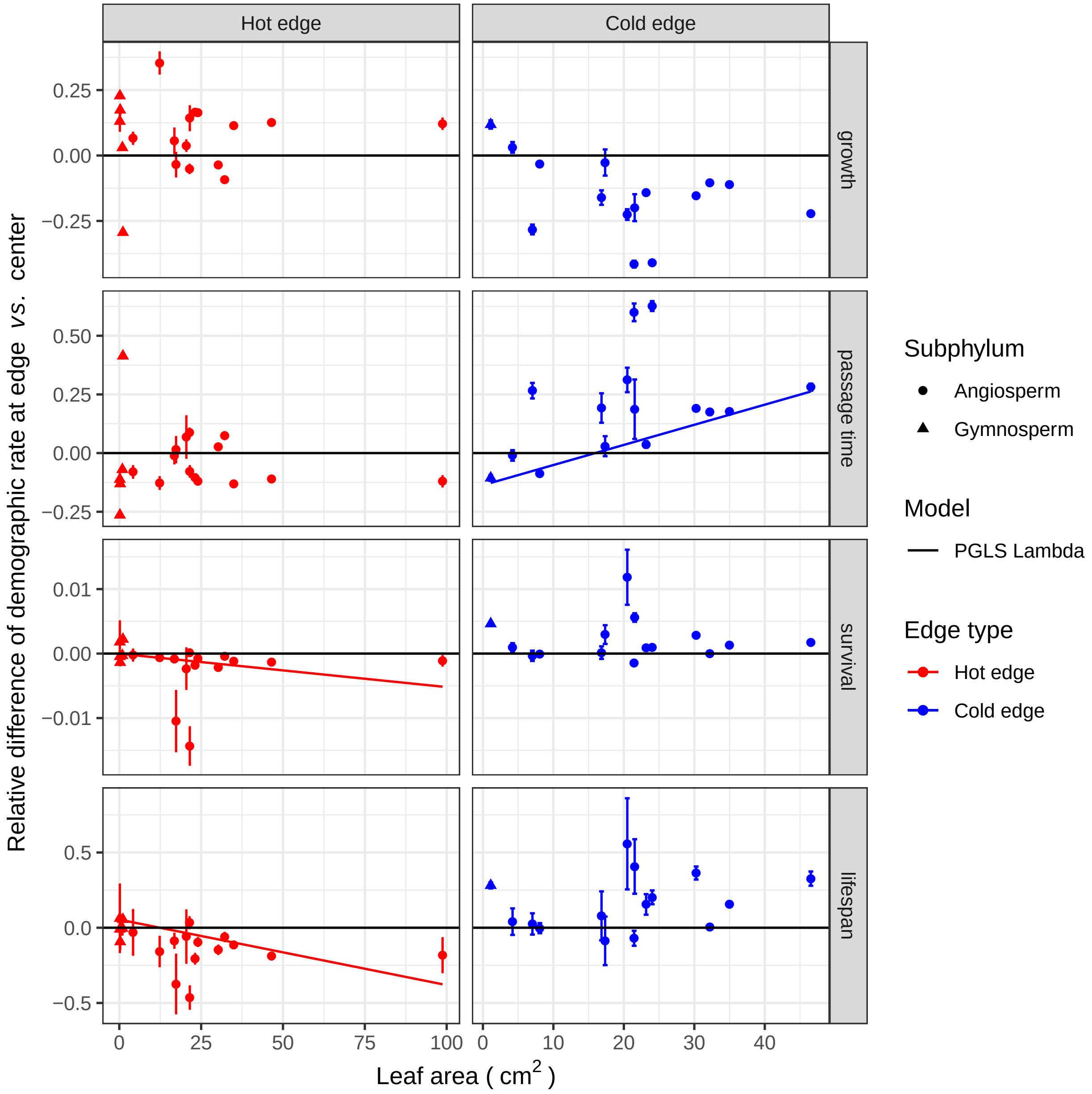

Figure 23: **Changes in demographic responses at the edge ($\Omega^m_{edge}$) in function of species leaf area.** Species demographic response at the edge - measured as the relative differences in the demographic metric at climatic edge *vs.* the median climate of the species distribution - as a function of species leaf nitrogen per mass. For each species the mean (point) and 95% quantiles (error bar) of the demographic response over the 100 data resampling is represented for both the hot (red) and the cold (blue) edges. Phylogenetic generalized least squares (PGLS) regressions are represented only for significant relationship with a non negligible magnitude of the effect. PGLS regression were first estimated by estimating the best value of Pagel's lambda (a measure of the phylogenetic signal), if the fit failed we fitted a PGLS model with a Brownian model corresponding to Pagel's lambda set at 1.

*Link between traits and species median climate*

Links between trait value and $\Omega^{m}_{edge}$ could either be related to leaf trait values resulting in better physiological tolerance and thus demographic performance at the edge, or to a correlation between the trait values and median climate of the species. In this last case, the relationship between the traits and the demographic response at the edge represents the effect of the median climate. Only wood density was significantly correlated to PC1 (Fig. 24).

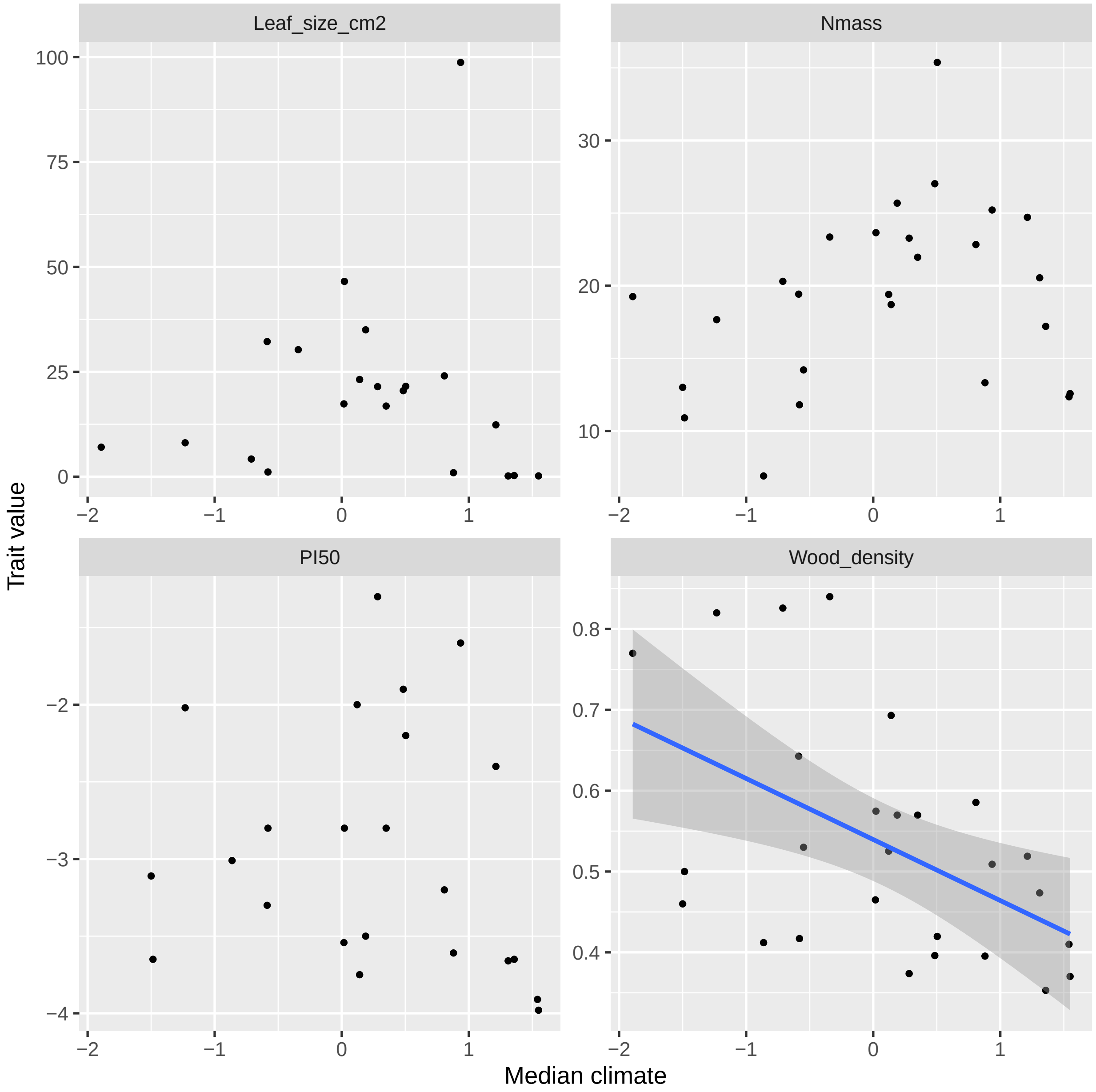

Figure 24: Correlation between species traits and species median climatic position one PCA axis one.

### COMPADRE elasticity analysis for vital rate of adult and juveniles

FunDivEUROPE NFI data cover only part of a tree's life cycle as they provide information only for tree larger than 10cm of dbh ('adult' part of the life cycle). To evaluate the contribution of the 'adult' part of the life cycle to population growth rate we used the COMPADRE database (Salguero-Gómez et al., 2015) that archives full matrix population models for plant to evaluate the contribution of the vital rates for the 'adult' and the 'juvenile' (less than 10cm of dbh) part of the life cycle to the population dynamics. There are too few species in common between COMPADRE and FunDivEUROPE to allow a species-level analysis, we thus extracted all tree species from COMPADRE divided them according to their shade-tolerance using Niinemets & Valladares (2006) (shade tolerant above 2.5 or below 2.5) as species with different shade-tolerances might have very different demographic strategies. Based on COMPADRE meta-data we extracted information on the size class that was closer to the 10cm dbh threshold to divide the matrix in 'juvenile' and 'adult'. Then we analysed the elasticity of the population growth rate to each vital rates of the 'juvenile' or 'adult' stage with a perturbation analysis of these vital rates. Elasticity gives the relative change in population growth rate for a relative change in vital rate.

We computed elasticity with the 'brut-force method' (Morris & Doak, 2002), by re-computing $\lambda$ after growth, survival, or fecundity of small (dbh < 10cm) or large tree (dbh > 10cm) was perturbed as $\frac{\lambda_o - \lambda_p}{v_o - v_p}$, where $v$ is the vital rate tested and the subscript $o$ and $p$ respectively represent the original or perturbed value.

#### *COMPADRE analysis results*

The analysis of the matrix population models extracted from the COMPADRE data base for 149 species shows that survival is one of the most important vital rate in term of population growth rate elasticity and this is the case for both small tree and large tree (see Fig. 25). This is inline with several analysis exploring the demographic signature of contrasted type of plant that show that survival is crucial for the population growth rate of trees (Adler et al., 2014). In contrast, growth had a smaller effect and mainly for small tree not covered by the FUNDIV data base. Fecundity, which is not monitored in the FUNDIV data base had an elasticity of the same range of magnitude as small tree growth (Fig. 25). Fig. 25 clearly shows that the demographic dynamics of tree larger than 10cm of dbh is crucial for population dynamics.

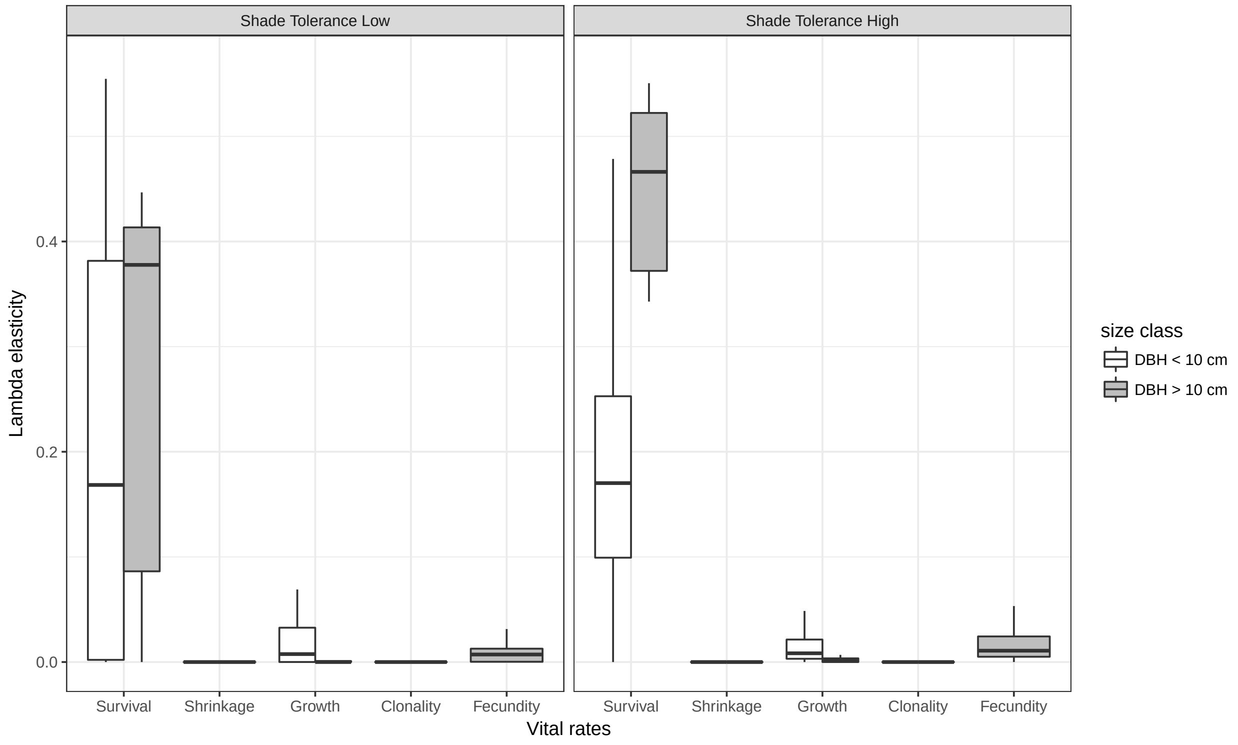

Figure 25: Elasticity of tree population growth rate for five vital rates and two size classes (below 10cm of dbh or above 10cm of dbh) for shade tolerant and shade intolerant tree species. Elasticity of lambda is computed from matrix population model of tree extracted from the COMPADRE open data base.
